# Transient transcription factor depletions explain diverse single-cell responses of LexA target promoters to mild DNA damage

**DOI:** 10.1101/2024.11.28.625836

**Authors:** Luca Galbusera, Gwendoline Bellement-Theroue, Thomas Julou, Erik van Nimwegen

## Abstract

In bacteria, the effects of transcription factors (TFs) on the expression of their target genes are highly stochastic at the single-cell level. Not only do TF concentrations fluctuate in time and from cell to cell, in each cell the actual binding and unbinding of TFs to promoters is also stochastic. However, how this ‘noise propagation’ from TFs to their targets determines the expression fluctuations in target genes of the same TF has so far not been quantitatively characterized. Here we use fluorescence time-lapse microscopy in combination with microfluidics and automated image analysis to quantitatively track the single-cell growth and gene expression dynamics of different target promoters of the LexA SOS response regulator in *E. coli* under mild DNA damage induced by the antibiotic ciprofloxacin (CIP).

We find that, at the single-cell level, LexA target promoters exhibit stochastic bursts in protein production with specific quantitative characteristics. First, the durations of protein production bursts are short, identical for all promoters, and independent of CIP levels. Second, the heights of the production peaks are exponentially distributed for each promoter, and independent of CIP level, but with different average heights for different promoters. Third, the frequency of the production bursts increases with CIP level for all promoters, but differs across promoters.

Importantly, we show that these observations are inconsistent with popular ‘equilibrium’ models of bacterial gene regulation that assume that the dynamics of binding and unbinding of TFs is fast relative to the time-scale of TF concentration fluctuations. Instead, all observations can be quantitatively fit by a relatively simple model in which DNA damage events lead to short transient dips in LexA concentration, and an expression burst occurs in a cell whenever LexA unbinds from the target promoter during such a short dip in LexA concentration. Moreover, because these transient dips in LexA concentration occur on the same time-scale as the stochastic unbinding of LexA from promoters, the response of a target promoter to an individual DNA damage event is stochastic and depends in a complex manner on the strengths of its LexA binding sites and the duration of the event. Consequently, the expression levels of different LexA target genes in a given cell reflect distinct statistics of the frequency and durations of DNA damage events in the recent history of that cell.

Our results show that, even in the simple scenario where a single TF is induced, different target promoters can exhibit distinct stochastic responses that depend on the precise kinetics of the induction events.

## Introduction

Bacterial cells can adapt their gene expression to grow in a large variety of environments and survive many different stresses. Much of this regulation occurs through the sequence-specific binding of transcription factors (TFs) to sites near the transcription start of genes, thereby altering the rate at which transcription is initiated [1–3]. Until about two decades ago most studies of gene regulation in bacteria concerned the bulk behavior in large population of cells. Since at the population level gene expression changes appear highly deterministic, this may have created the impression that gene regulation is deterministic and maybe even finely tuned. However, since genes are typically only present in one or two molecular copies per bacterial cell, TF binding and transcription initiation are single molecule events and are therefore subject to inherent thermal molecular noise. That is, basic biophysical principles predict that gene expression should exhibit substantial random fluctuations at the single cell level. Indeed, early studies of gene expression at the single cell level not only showed that gene expression varies between identical promoters in the same cell, but also that the actions of TFs can propagate noise so as to increase the cell-to-cell variation in gene expression of their targets [4, 5].

Recent work from our lab has shown that noise propagation from regulators to their targets is a key determinant of gene expression noise on a genome-wide scale in *E. coli*, that highly regulated promoters tend to be the most noisy, and that gene expression noise is highly condition-dependent [6, 7]. This raises the question of how noise is propagated from regulators to their targets, i.e. how the actions of TFs determine the gene expression dynamics of their targets in single cells.

A popular modeling approach is to assume that both the dynamics of binding and unbinding of TFs to their target sites and the transcription initiation events are fast relative to fluctuations in TF concentrations [8–10]. Under that assumption, the fractions of time that different TF binding configurations occur at a promoter can be calculated as a function of the TF concentrations assuming effective ‘thermodynamic equilibrium’ in the TF binding dynamics. Each of the possible TF binding configurations at a promoter is associated with a rate of transcription initiation, and the expected output from a promoter can then be calculated by averaging over the fractions of time the promoter is in each binding configuration. In this ‘equilibrium’ model, the output from each promoter is a deterministic function of the TF concentrations in the cell, and any noise propagation from regulators to their targets is entirely due to cell-to-cell fluctuations in TF concentrations.

However, it is currently unclear to what extent these assumptions actually hold for bacterial regulatory circuitry. For example, it cannot be excluded that fluctuations in TF concentrations can be sufficiently fast that the equilibrium assumption breaks down. More generally, while equilibrium models predict the expected fractions of time a given promoter will be in different TF binding configurations, the inherent stochasticity in the binding/unbinding dynamics may cause the actual fractions of time a promoter spends in different configurations to fluctuate from cell-to-cell, and this may also drive variations in gene expression.

To investigate how regulators affect the expression dynamics of their targets in single cells we decided to focus on the LexA regulon in *E. coli*, which is activated in response to DNA damage [11, 12]. This regulatory system presents several characteristics that make it well suited to quantitatively study noise propagation at the single cell level. First, the system has been intensively studied, its induction mechanism has been dissected in substantial detail, and its target genes have been well characterized, including the sequence specificity of LexA and its binding sites in target promoters. Moreover, many of the parameters of this system such as LexA protein levels and the unbinding rates from different LexA target sites, have been determined with reasonable accuracy, and this strongly constrains possible models of the regulatory dynamics of this system. In addition, the regulon can be easily induced in the laboratory by damaging DNA either through UV light or using particular antibiotics.

Briefly, the SOS pathway comprises genes involved in the repair of DNA damage and is regulated by the repressor LexA which, in the absence of DNA damage, is bound to the promoters of its target genes, keeping them repressed. When DNA damage occurs, the protein RecA will form filaments on single-stranded DNA and LexA molecules auto-cleave on these RecA filaments, leading to a decrease in LexA concentration. This decrease in LexA concentration then leads to induction of its target genes, which include LexA and RecA themselves. Once the DNA damage is repaired, the RecA filaments disassemble, and the LexA expression recovers [11–15].

So far the induction of the LexA regulon has been studied mostly in response to UV radiation, and initially at the bulk population level, leading to several models for the regulatory dynamics at the population level, e.g. [15–18]. However, when the induction dynamics in response to UV radiation was measured at the single cell level, it became clear that the population measurements mask a more complex single-cell dynamics with multiple roughly synchronized peaks in the activity of LexA target promoters [19]. However, because UV radiation induces a complex cascade of downstream effects including major physiological changes such as filamentation and growth arrest, it is challenging to disentangle how the various processes downstream of UV radiation cause the observed stochastic peaks in LexA target expression. Indeed, a host of different models has been proposed in the literature to explain these complex temporal modulations in activity of LexA target promoters [17, 20–22].

We thus reasoned that expression regulation of LexA targets in single cells might be easier to interpret in a regime where there is only mild DNA damage and induction of the SOS response. Recent time-lapse microscopy studies with a fluorescent transcriptional reporter of the LexA target promoter RecA have shown that, even in the absence of stressors that induce DNA damage, there are stochastic epxression bursts of this promoter in single cells [23, 24], suggesting that stochastic induction of LexA targets in single cells can be readily observed even at low levels of DNA damage.

Our approach to understanding the single cell gene regulation of the LexA regulon was as follows. We first selected a set of genes that have been identified as targets of LexA and that are currently believed not to be regulated by any other TF. Using fluorescent transcriptional reporters of the promoters of these target genes [25] we tracked growth and gene expression of these target genes in lineages of single cells using an integrated setup that combines fluorescence time-lapse microscopy with microfluidics and automated image analysis [26, 27]. Using the DNA damaging antibiotic Ciprofloxacin (Cipro) [28] we exposed the cells to increasing levels of mild DNA damage, increasingly inducing the LexA targets without substantially affecting either the growth rates or the physiology of the cells, i.e. avoiding filamentation. Using this setup, we quantified the time-dependent stochastic gene expression for different LexA target promoters as a function of the concentration of Cipro.

Below we first show that, at the population level, different promoters show different levels of induction as a function of Cipro, and then show that at the single-cell level, the induction is driven by short stochastic bursts in expression. We then show that the frequencies, heights and durations of these gene expression bursts have very specific quantitative characteristics and that ‘equilibrium’ models of gene regulation are unable to reproduce these observed characteristics. Instead, we show that all our data can be accurately fit by a model in which stochastic DNA damage events cause short transient dips in LexA levels and that expression bursts occur whenever LexA unbinds from the promoter during such a transient dip. In the final result section we use simulations of this model to show that, because the durations of the dips in LexA concentration are comparable to the time for LexA to unbind from its bindings sites, the response from each promoter is a complex stochastic function of the frequency and durations of DNA damage events in the recent history of the cell. Our results show that, even for the relatively simple example of mild induction of the LexA regulon by DNA damage events, different target promoters show highly distinct responses to the same induction events, which depends in a complex manner on the detailed time dynamics of the inducing signal.

## Materials and Methods

### Strains and transcriptional reporters

We measured fluorescence in single cells for a set of *Escherichia coli* MG1655 strains that each carry a transcriptional reporter inserted upstream of a gene for the fluorescent protein GFP-mut2 and expressed from a low copy-number plasmid [25]. All the sequences were verified by Sanger sequencing. We chose reporters corresponding to promoters that are known to be regulated by LexA and currently believed to not be targeted by any other TF [29]: ftsK, lexA, polB, recA, recN, ruvA, umuD, and uvrD. In addition, as a control we also measured fluorescence from analogous reporters of two synthetic promoters that were obtained by screening a large library of random sequence fragments for driving expression at either the median or at 97.5^*th*^ percentile of all native *E. coli* promoters [6]. Throughout the paper, we refer to these two synthetic promoters as Synthetic Medium and Synthetic High.

### Time lapse microscopy

Mother Machine experiments were performed over 30 hours as described in [26], using M9 + 0.4% glucose. One phase contrast and one GFP fluorescence image (Lumencor Spectra X, Cyan LED at 17% with ND4 filter, 2000 ms exposure) were acquired over multiple positions every 3 minutes. In order to study three increasing concentrations of Ciprofloxacin (Sigma Aldrich) consecutively, we used two media inputs, without Cipro and with 2ng/mL respectively, and set the flow rates so as to expose the bacteria to the following ratio of those inputs: 100/0 (first six hours, no Cipro), 50/50 (from 6h to 18h, 1ng/mL), 0/100 (from 18h to 30h: 2ng/mL). We refer to [26] for a detailed description of the Dual Input Mother Machine design that allows accurate dynamic control of the growth media that flow to the growth channels. To avoid perturbing the response to Cipro during the experiment, the selection antibiotic for the reporter plasmid (50*µg* / mL of kanamycin) was supplemented during the overnight preculture only. We note that plasmid loss during the experiment would lead to a gradual decrease of the fluorescence level in the corresponding cell due to cytoplasm dilution, eventually producing non-fluorescent cells. Remarkably, we do not observe any cell becoming non fluorescent during the 30 h experiments, which indicates a very robust partitioning mechanism for the pSC101 origin. All experiments were performed at 37°C.

### Data preprocessing

To measure cell sizes and their gene expression, we analyzed the images using the Mother Machine Analyzer (MoMA) software as described in [26], which provides accurate segmentation and tracking of the cells.

From the phase contrast imaging data we obtain estimates of the cell length at each time point and from the fluorescence data we obtain estimates of the total fluorescence using a method described previously [26]. Briefly, the fluorescence signal across the channel shows a Cauchy-shaped peak that decays to background fluorescence between the channels, and the amplitude of this peak is proportional to the total number of GFP molecules in the cell. We previously estimated the conversion factor between the fluorescence peak amplitude and GFP molecule number from the fluctuations in the distribution of fluorescence at division [26] and we use this to convert fluoresence intensities into estimated GFP molecule numbers.

### Analysis of the data

We find that the estimated number of GFP molecules generally correlates strongly with the size of the cell (Suppl. Fig. S1), especially for the constitutive synthetic promoters. We thus decided to measure gene expression as GFP concentration, which we define as estimated total number of GFP molecules divided by estimated cell length (which is proportional to cell volume since cells generally keep their width constant in these growth conditions).

Since GFP concentration changes smoothly in time, measurements of GFP concentration from consecutive time points of the same cell are highly correlated. Therefore, to calculate distributions of GFP concentrations across the ‘population’ of cells, we use averages over time segments of length roughly equal to the correlation time of GFP concentration fluctuations.

As explained in section 1.2 of the Supplementary Material, we observed some day-to-day variability in the absolute fluorescence levels of the same constitutive synthetic promoter, which are likely due to subtle uncontrolled variations in the growth-conditions across days (Suppl. Fig. S2). In contrast, we find that relative expression levels are highly reproducible across days. To compare results from different days we thus normalized expression levels from different days by the average expression of the synthetic high promoter measured on that day (see Suppl. Mat. section 1.2 and Suppl. Fig. S2).

Finally, we noticed that due to the effects of the illumination, cell behavior transiently adapts during the first 1-2 hours of the experiment, and we therefore discarded all measurements taken during the first two hours (Suppl. Fig. S3 and Suppl. Mat. section 1.3).

### Estimating volumic GFP production rate

We observed that, in general, the rate of GFP production is proportional to cell volume *V*, which is consistent with an approximately constant concentration of ribosomes in the cells leading to an overall production of proteins proportional to cell volume. As in previous works from other labs, e.g. [19, 24], we thus write the time-dependent total production rate of GFP as a product of cell size *V* (*t*) and a volumic production rate *q*(*t*), which we use to quantify the activity of the promoter. Further, the total GFP fluorescence of a cell decays at a constant rate *β* due to the combined effects of bleaching and dilution. We thus obtain the following differential equation for the total GFP fluorescence *G*(*t*) of an exponentially growing cell

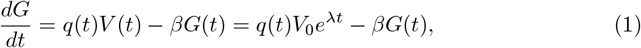

where in the second equation we have assumed that the volume *V* (*t*) grows exponentially at a rate *λ* starting from a volume *V*_0_ at the beginning of the cell cycle. We have previously used control experiments to measure *β* [26] and, as detailed in Suppl. Mat. section 2, we combine estimates of the instantaneous growth-rate *λ* from the phase contrast measurements with measurements of the total fluorescence *G*(*t*) to estimate the instantaneous production rate *q*(*t*) during short time windows using the above equation. In particular, to obtain a local estimate for the volumic production rate *q*(*t*) at time *t*, we solve the above differential equations for a time window centered on *t* spanning 5 acquisition time points (that is, covering 15 minutes) and estimate the local growth rate using a window of 9 time points centered on the same time point (since, empirically, we find that growth rate varies relatively little over this 27 minute time scale). We checked that reducing the width of these windows has no significant effect on the statistics of peaks in *q*(*t*), apart from slightly reducing the peak widths.

### Peak identification

The GFP production traces of several LexA target promoters show clear bursts, i.e. relatively short-lived peaks in volumic production *q*(*t*). These peaks in volumic production are characterized by exponential tails in the distributions of *q* (Suppl. Fig. S4). To characterize these bursts, we defined production peaks as any contiguous segment of the volumic GFP production *q*(*t*) above a given cut-off *q*_*c*_, and we fit *q*(*t*) in each such segment to a Gaussian curve from which we measure the height and duration of the peak. In Suppl. Mat. section 3 we give a detailed description of our peak-annotation algorithm, and show that the statistics of the peaks are insensitive to the precise value of the cut-off used (Suppl. Figs. S5, S6, S7, and S8).

## Results

### LexA target genes show different levels of induction at the population level

To study the induction of the LexA regulon we selected 8 promoters that are known to be regulated by LexA and are currently believed not to be regulated by any other TF (apart from the sigma factor Sigma70) [29]. In addition, we included two synthetic promoters that we previously obtained by screening a library of completely random short sequence fragments for fragments that drive expression [6]. These promoters, which we denote by ‘synthetic high’ and ‘synthetic medium’, exhibit average expression levels at the 97.5*th* and 50*th* percentile of all native *E. coli* promoters, respectively. In addition, these synthetic promoters do not appear to contain TF binding sites other than a Sigma70 site [6] and were observed to be constitutively expressed across a range of growth conditions [7]. We thus use these synthetic promoters as constitutively expressed controls.

We induced the LexA target promoters using low levels of the DNA damaging antibiotic Ciprofloxacin. Our microfluidic setup enables controlled variation of the Cipro concentration over time, allowing direct observation of gene regulatory responses to different induction levels in single cells [26]. In particular, we monitored the cells over a period of 30 hours, starting with no antibiotic in the first 6 hours, switching to 1 ng/mL in the following 12 hours and finally to 2 ng/mL in the final 12 hours. Note that these concentrations are all far below the MIC of Ciprofloxacin and do not cause major physiological changes to the cells such as filamentation. Indeed, at these concentrations the growth rates are not noticeably affected by the presence of the antibiotic (Suppl. Fig. S9).

In Fig 1A, we show snapshot images across time of cells and their fluorescence levels from one growth lane containing cells that express GFP from the synthetic high promoter (top) and one growth lane containing cells expressing GFP from the recA promoter (bottom). We can see that, while the synthetic promoters exhibit approximately constant expression across the different Cipro concentrations, recA promoters exhibit much more variable expression which on average increases as the concentration of Cipro increases. This is shown more quantitatively in Fig 1B, which shows the fold change of the mean fluorescence level (averaged over all cells) as a function of time for the synthetic high promoters and the 5 LexA target promoters that were induced in our setup. Note that we did not observe a detectable induction for 3 additional promoters (ruvA, uvrD, and umuD) that are reported to be regulated by LexA (Suppl. Fig. S10).

**Fig 1.**
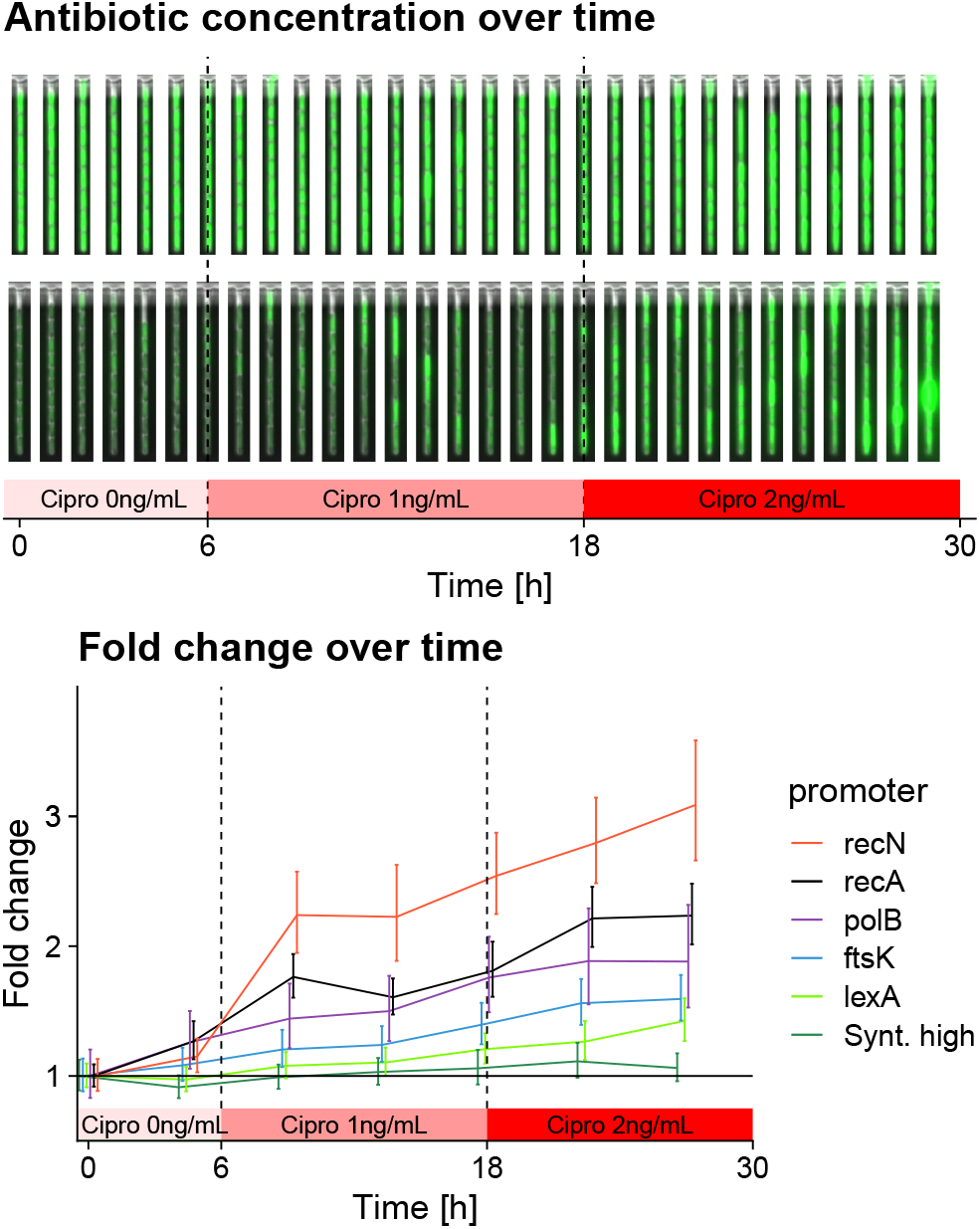
Induction of promoters under control of LexA. **A**: Time-lapse of *E. coli* growing under increasing subinhibitory concentrations of Cipro. The upper strip shows cells expressing GFP from the synthetic high promoter, while the bottom strip shows cells expressing GFP from the recA promoter. Note that Cipro is known to induce DNA strand breaks. **B**: Fold-changes in average expression as a function of time for 5 different LexA target promoters, and the unregulated synthetic high promoter (mean± one standard-error).

Importantly, Fig 1B shows that the expression of different promoters is upregulated to different extents in response to the same change in conditions. In addition, Suppl. Fig. S10 shows the basal GFP concentrations (in the absence of Cipro) and the fold changes for all the LexA regulated promoters.

### GFP production from LexA regulated promoters occurs in short bursts

To better understand how different target promoters respond to the same induction of the LexA TF, we monitored the expression dynamics in single cell lineages. For each cell, the microfluidic setup enables us to track the total GFP intensity over time, as shown in the top panel of Fig 2A. To quantify gene expression from each promoter we define the volumic GFP production rate *q*(*t*) by modeling the dynamics of the cells’ total GFP fluorescence *G*(*t*) as a function of time *t* using the following equation:

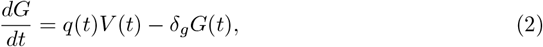

where *q*(*t*) is the volumic production rate, *V* (*t*) the volume of the cell, and the last term represents the loss in fluorescence due to bleaching and GFP protein decay, which occurs at rate *δ*_*g*_.

**Fig 2.**
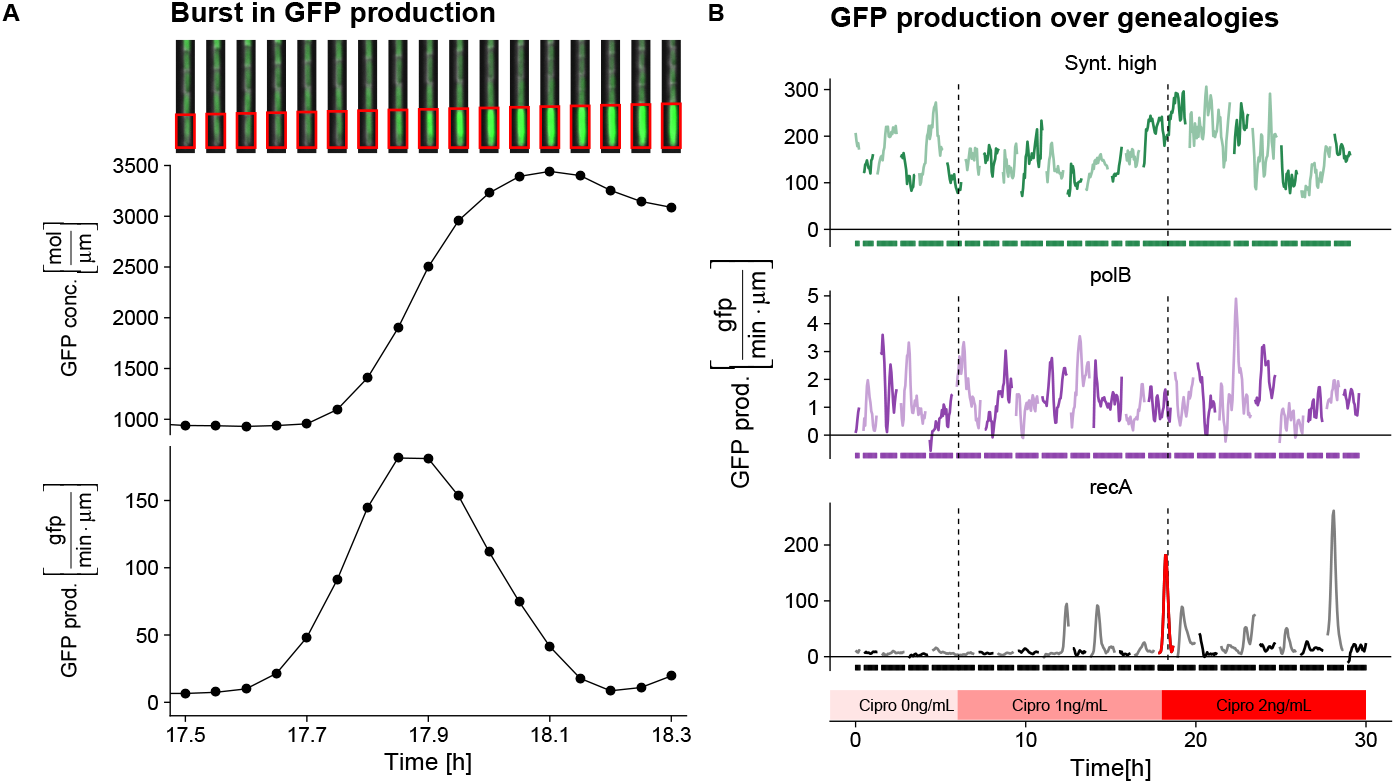
GFP production over time for single cells. **A**: Time-lapse images highlighting a sudden increase in GFP fluorescence for a cell expressing GFP from the recA promoter (top, red rectangles), the corresponding GFP concentration (middle) and inferred volumic GFP production *q*(*t*) (bottom). The production shows a transient burst from 17.7 to 18.1 hours (approximately 20 minutes). **B**: Example of volumic GFP production *q*(*t*) along the entire duration of the experiment for three promoters. For each promoter, the production *q*(*t*) of only one representative cell genealogy is displayed, and the boundaries between the cells of the genealogy are shown by the rectangles below each trace. The dotted vertical lines separate the three different Cipro concentrations. The red cell in the recA trace corresponds to the cell shown in panel A.

Although our analysis does not depend on a specific biophysical interpretation of the volumic production rate *q*(*t*), note that if the concentration of active ribosomes and the average translation rate of all mRNAs in the cell are both roughly constant across the cell cycle, then *q*(*t*) is proportional to the fraction of GFP mRNAs among all mRNAs in the cell at time *t*. Since mRNAs turn over on a time scale of just a few minutes [30], the production rate *q*(*t*) thus corresponds to the recent transcriptional activity of the promoter. As detailed in the Supplementary Material section 2, we estimate *q*(*t*) from measurements of total fluorescence in short sliding windows, assuming that *q*(*t*) changes smoothly within these windows.

We observed that the traces of volumic production *q*(*t*) across time for LexA target promoters exhibit relatively short bursts. As an example, Fig. 2A shows the raw fluorescence images, GFP concentration, and volumic production in a single cell carrying the GFP reporter of the recA promoter during a 45 minute time segment, illustrating the occurrence of a production burst. Examples of time traces of GFP production over the entire experiment are shown in Fig 2B for three representative lineages of cells expressing GFP from different promoters. There is a clear difference between the time traces of the LexA targets and that of the constitutively expressed promoter. Whereas the production of the synthetic high promoter fluctuates in a manner resembling a random walk around a mean value, the LexA target promoters recA and polB are characterized by a low background production interspersed by production bursts, which are especially clear for the recA promoter.

### Characterization of the expression bursts

Although the expression bursts are very clear for some promoters, e.g. recA, for others such as polB, it is more challenging to distinguish the bursts in production from the ‘background expression’. To more rigorously characterize and quantify the production bursts, we calculated the distribution of volumic production rate *q* across all cell lineages for each promoter and observed that, in contrast to the constitutive promoters, all LexA targets exhibit exponential tails in the distribution of *q* (Suppl. Fig S4). In addition, we further verified that *q* values in these tails correspond to peaks in the time traces of volumic production. As detailed in the Supplementary Material section 3, we define bursts in production as local maxima in the time traces of volumic production *q*(*t*), whose heights lie within the exponential tail of the distribution of *q*. In addition, we estimate the height and width of each production peak by fitting the trace *q*(*t*) around the local maximum to a Gaussian function. In particular, we define the width of the peak as the standard-deviation of the fitted Gaussian.

Figure 3 shows the distributions of the heights and widths of production peaks that we find for different promoters and at different levels of Cipro. For clarity, we here only show the distributions for lexA, recA, and recN and show analogous results for polB and ftsK in Suppl. Fig. S6 and S7. These supplementary figures additionally show that the peak statistics are insensitive to the cut-off used to define the peaks.

**Fig 3.**
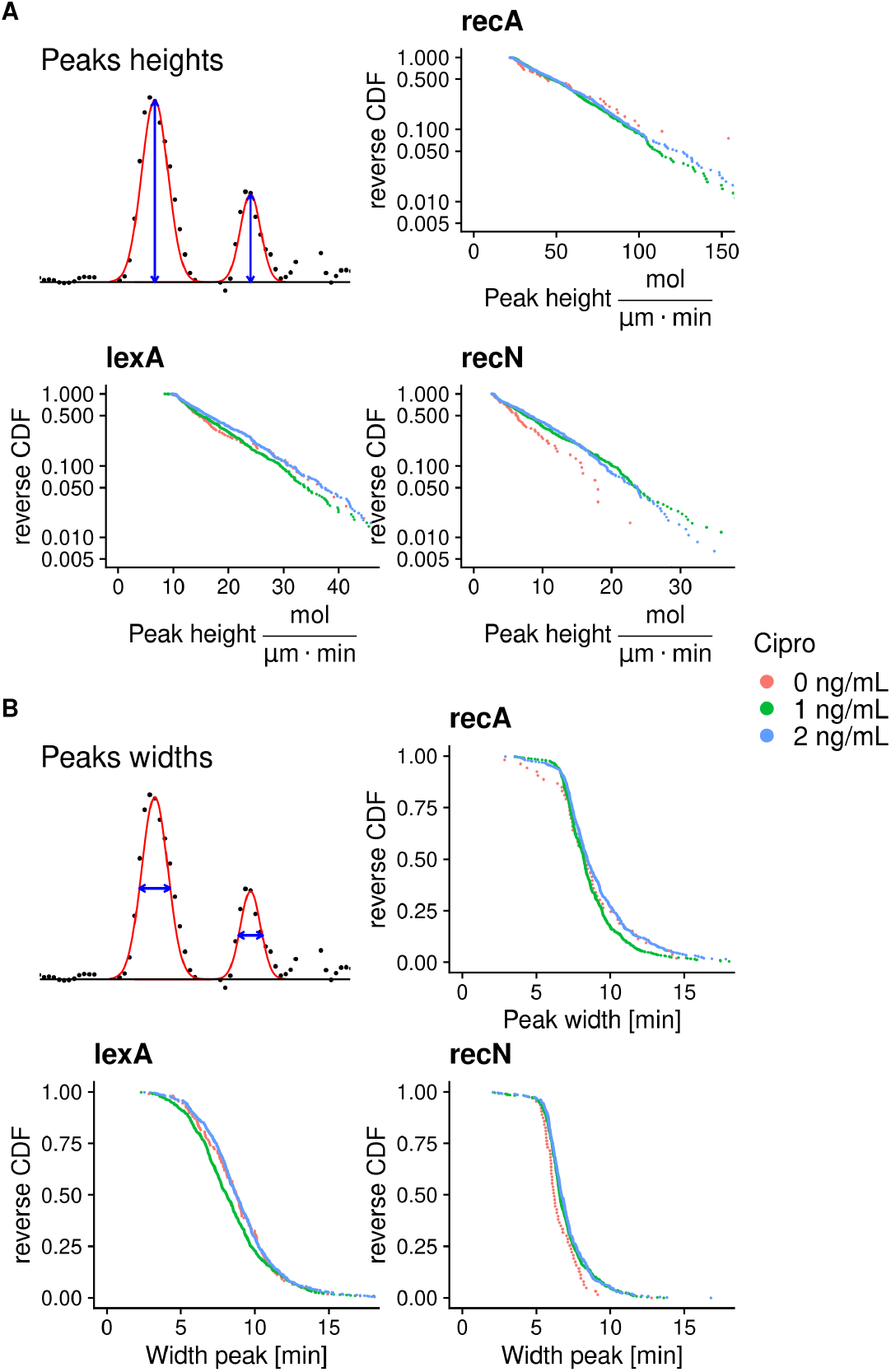
Quantitative characteristics of the bursts in volumic GFP production. **A**: Bursts in volumic production are fitted to Gaussian shaped peaks to estimate their heights (top left panel). The other panels show reverse cumulative distributions of the peak heights for three target promoters (different panels) and for the three Cipro concentrations (colors, see legend). Note that the vertical axes are shown on logarithmic scales. **B**: Reverse cumulative distributions of peak widths for the same three promoters and Cipro concentrations. Note that peak heights are exponentially distributed and independent of the Cipro concentration, but with a different mean for each promoter. Peak durations have the same distribution across all promoters and Cipro concentrations.

First, we observed that the heights of the production peaks are almost perfectly exponentially distributed and independent of Cipro concentration for each promoter (Fig. 3A). That is, each promoter has its own exponential distribution of peak heights, with a different average peak height, that is unchanged by changes in the Cipro concentration. Second, we observed that the distributions of peak durations are not only independent of Cipro concentration, but also virtually identical for all three promoters, with a median duration of around 8 minutes (Fig 3B). Note that, since the peak duration is defined as the standard-deviation of the fitted Gaussian, the entire duration from the start to the end of the production burst is about 20 − 30 minutes.

Finally, to estimate the relative frequencies of bursts across promoters and Cipro concentrations, we extend the exponential tails of the peak frequency as a function of the height cut-off back to zero, and in this way estimate an effective ‘pseudo-frequency’ of each promoter at each Cipro concentration (see Suppl. Material section 3.4). We find that these pseudo-frequencies increase with the concentration of Cipro in a similar although not strictly identical manner across promoters (Fig 4).

**Fig 4.**
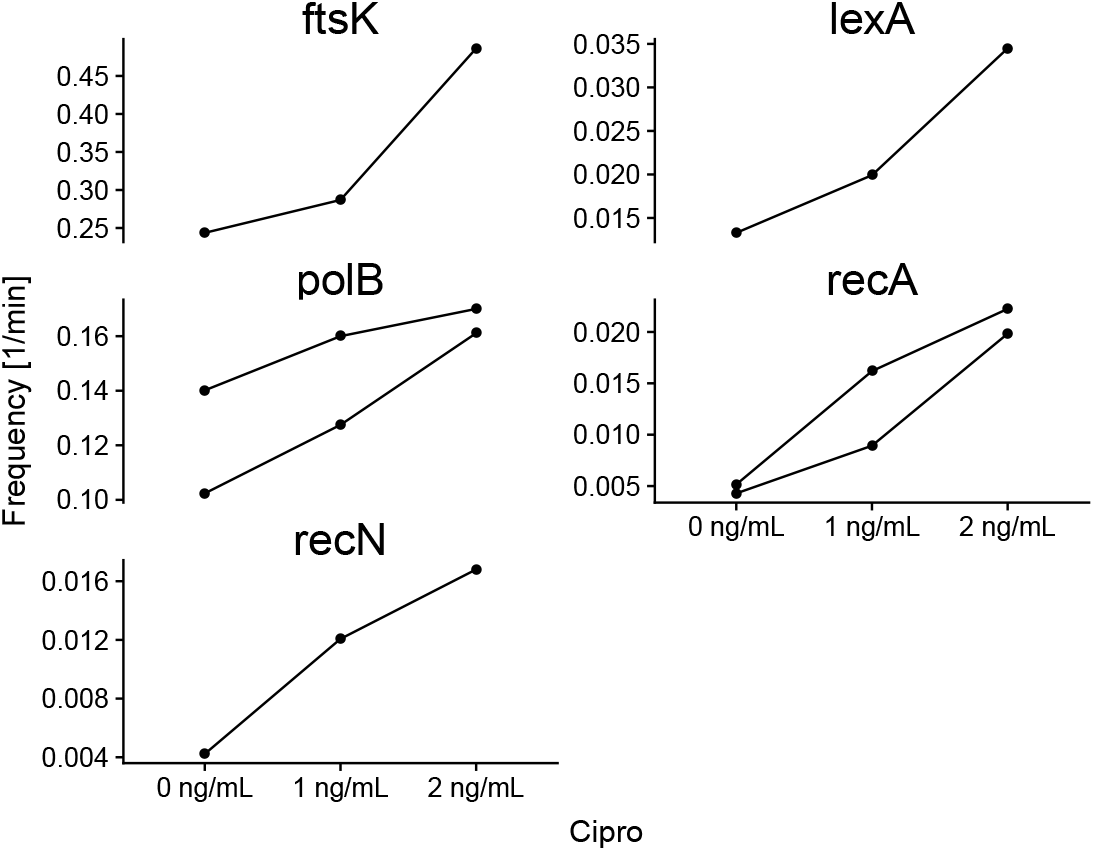
Relative frequencies of the bursts in GFP production as a function of Cipro concentration across LexA target promoters. Each panel shows the estimated pseudo-frequency of production bursts (see text) as a function of Cipro concentration for one LexA target promoter (indicated at the top). Note that the two curves for polB and recA show results from measurements on two separate days. The burst frequency increases monotonically with Cipro concentration for all promoters but rises more steeply for some promoters (i.e. recN) than for others (ie. polB).

In summary, we find that all target promoters exhibit production bursts which become more frequent as the Cipro concentration increases, but whose durations are identical for all promoters, and whose heights depend on the promoter, but are independent of Cipro concentration. These quantitative observations put strong constraints on the possible biophysical mechanisms underlying these production bursts, as we explore in the next section.

### A simple model of stochastic single-cell gene expression

To explore what single cell regulatory mechanisms can explain our observations we will use a stochastic modeling framework that explicitly incorporates cell growth, binding and unbinding of LexA to the promoter, transcription events, translation events, and so on. In particular, the model follows a single cell lineage through time including the exponential growth of the cell and division events. Following the observation that added length between birth and division is approximately independent of growth-rate and cell size at birth [31, 32] we model the cell cycle by, after each division, drawing a random added length Δ*L* from a Gaussian distribution with mean and variance matching those observed in the experimental data. We then let the cell grow exponentially until a length Δ*L* has been added, upon which it divides again. For simplicity, we ignore growth-rate fluctuations and let all cells grow at the same rate, given by the median observed growth rate in the data (doubling roughly once every 80 minutes). Our model also ignores small cell size asymmetries at cell division and assumes the mother cells divide exactly in half at each cell division.

Our model of the gene expression dynamics of a LexA target promoter in a given cell includes the stochastic reactions illustrated in Fig. 5A. We assume that the promoter can exist in either a LexA-bound or LexA-free state and that it switches stochastically between these two states. LexA unbinds at a rate *L*_off_, which depends on the affinity of the binding site(s) in the promoter, and LexA binds at a rate *L*_on_ which we assume is proportional to the concentration of free LexA in the cytoplasm.

**Fig 5.**
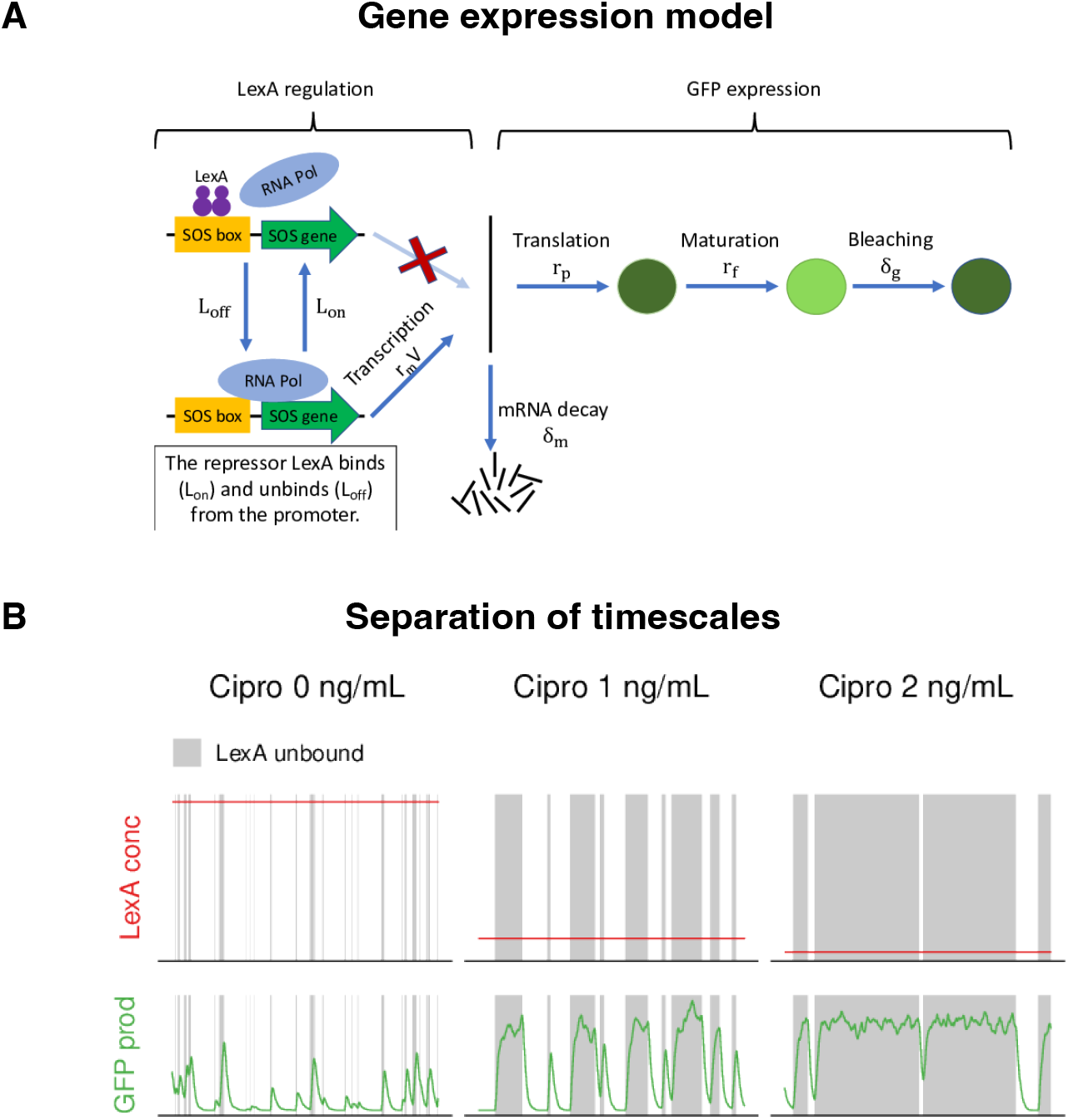
Gene expression and equilibrium regulation models for the LexA regulatory network. **A**: Our gene expression model for a LexA target promoter includes stochastic switches between a transcribing state (unbound) and a repressed state (bound by LexA) as well as stochastic transcription events, translation events, mRNA decay events, GFP maturation, and bleaching/decay, each occurring at a given rate per molecule. **B**: Example simulation of the ‘equilibrium’ model where the concentration of LexA is assumed to be effectively constant over time and decreasing with the Cipro concentration (top row). In this model, expression bursts correspond to periods where the promoter is unbound (gray intervals), leading to peaks in volumic production (bottom row). Note that, as the average concentration of LexA decreases, LexA takes longer to rebind, causing production bursts to increase in duration.

In general, LexA target promoters will transcribe at a lower rate when bound by LexA than when unbound. To reduce the number of free parameters, we make the idealization that promoters only transcribe when unbound, i.e. the transcription rate in the bound state is zero. In the unbound state, we assume transcription events occur at a rate *r*_*m*_*V*, i.e. proportional to the volume *V* of the cell. That is, we do not explicitly model the dynamics at each independent copy of the plasmid (which we previously determined is tightly regulated to roughly a handful of copies per chromosome [7]) but simply assume the transcription rate scales with cell volume as plasmid copy number grows with cell volume.

Each mRNA has a probability *δ*_*m*_ per unit time to decay and a probability *r*_*p*_ per unit time to be translated into a GFP molecule. In order to correctly capture the time scales of the bursts, we also take into account the maturation of the GFP molecules. Each GFP produced is initially immature (i.e. non-fluorescent) and has a probability *r*_*g*_ per unit time to mature into a fluorescent protein. Finally, each GFP has a probability *δ*_*g*_ per unit time to decay, which represents the combined effects of photo-bleaching and protein decay [26]. Note that many of the parameters of this model, such as the mRNA decay rate, the translation rate of the GFP reporter, the GFP maturation rate, and the rate of LexA unbinding from the RecA and consensus binding site are known from the literature, at least approximately. We used this knowledge to set many of the parameter values of the model (see Table 1).

**Table 1.** Parameters of the gene expression model. Green cells correspond to parameters for which an approximate value is known from the literature. For orange cells a range of possible values is fixed by the requirements of fitting to our observations (see Suppl. Mat. section 6).

| Rate | Description | Value / Constraint |
| --- | --- | --- |
| $\delta_g$ | GFP bleaching+decay rate | $0.0032 \text{ min}^{-1}$ [26] |
| $r_g$ | GFP maturation rate | $1/8 - 1/6 \text{ min}^{-1}$ [53, 54] |
| $r_p$ | Translation initiation rate | $20 \text{ min}^{-1}$ [55–58] |
| $\alpha$ | LexA decay rate during DNA damage | $1.5 \text{ min}^{-1}$ [48] |
| $\beta$ | LexA recovery rate after repair | $1.5 \text{ min}^{-1}$ |
| $L_{ss}$ | steady-state LexA binding rate | $200 \text{ min}^{-1}$ |
| $L_{off}$ | LexA unbinding rate | $0.1 \text{ min}^{-1}$ (consensus sequence),<br>$3 \text{ min}^{-1}$ (recA site) [18, 59, 60] |
| $\delta_m$ | mRNA decay rate | Between $1/2 - 1/6 \text{ min}^{-1}$ [61] |
| $r_r$ | DNA damage repair rate | $0.5 - 1.0 \text{ min}^{-1}$ |
| $r_m$ | transcription initiation rate per unit volume for the unbound promoter. | Fixed by $r_r$ and observed average peak height. |
| $r_b$ | Frequency of DNA damage events | Fixed by observed peak frequency. |

The final quantity to specify in this model is the dynamics of LexA concentration in the cell and its dependence on the concentration of the antibiotic Cipro. Below we will first explore a model in which there is an effectively constant LexA concentration at each antibiotic level and show that this leads to predictions that are clearly at odds with our observations.

### The observed burst characteristics disagree with predictions of ‘equilibrium’ models

As mentioned in the introduction, one of the assumptions most frequently adopted in modeling gene expression in bacteria assumes that binding configurations at promoters are in statistical equilibrium given TF concentrations, e.g. [33]. Indeed, starting from the seminal work of Ackers et al. [34], this strategy has been applied in a large number of studies, including systems with relatively complex regulation, e.g. [8, 35–42].

The key assumption of these ‘equilibrium’ models is that the production from each promoter is an average of transcription rates in different binding configurations, which assumes that the time scales of TF binding and unbinding from the promoters, as well as the time scale of transcription initiation events, are fast relative to the time scale of LexA concentration changes [10, 33].

To investigate the burst characteristics predicted by the equilibrium assumption, we considered the idealized scenario in which, during each time segment at a given Cipro concentration, the promoters are experiencing an effectively constant LexA concentration, which decreases with Cipro levels (Fig 5B, top row). We used our gene expression model to investigate this parameter regime and representative results of simulations of the model are shown in Fig 5B. In this regime, the bursts in production correspond to periods of time when LexA is unbound from the promoter. As the concentration of Cipro increases, and the average concentration of LexA decreases, the rate at which LexA binds to unbound promoters is lowered, causing the time intervals in which the promoter is free from LexA to increase in duration (Fig 5B, top row). This in turn causes higher production bursts of longer durations (Fig 5B bottom row).

These predictions of the model clearly contradict the experimental observations of Figs 3 and 4, which show peak heights and durations are independent of Cipro concentration, whereas their frequency increases with Cipro concentration.

One could argue that our assumption of constant LexA levels is too restrictive, and that the equilibrium model allows LexA levels to vary in time and across cells, as long as these variations are slow relative to the rates of LexA binding and unbinding to the promoter. However, unbinding rates of LexA binding sites have been measured and can be as low as once every several minutes [18]. Therefore, in order for the equilibrium assumption to hold, LexA concentration changes must be slow relative to this time scale, and since production bursts last on the order of 10 − 20 minutes, this in turn implies LexA concentration must be roughly constant on the time scale of the production bursts. Furthermore, since bursts correspond to periods in which the promoter is unbound, and the rebounding rate is set by the LexA concentration, in the equilibrium model the distribution of burst heights and durations must reflect the distribution of LexA concentrations within a time segment. Therefore, as the LexA levels decrease with increasing Cipro concentration, the equilibrium model necessarily predicts longer and higher bursts, which contradicts the experimental observations. Therefore, it is clear that explaining the experimental observations requires a model in which LexA concentration fluctuations must occur on the same time scale as the observed bursts in production.

### Production bursts correspond to stochastic transient dips in LexA concentration

A successful model needs to explain why the durations of production bursts are independent of promoter and Cipro concentration, and why peaks heights are exponentially distributed and also independent of Cipro concentration. We start by considering what is known about LexA concentrations in the cell, and the time-scales for its binding and unbinding from target sites in promoters. The time-scale for LexA to unbind from one of its sites ranges from just seconds for weak binding sites to almost 10 minutes for the consensus site [18]. This already suggests that not every LexA unbinding event can generate a production burst, because production bursts are observed much less frequently.

Next is the LexA binding rate in conditions without DNA damage, i.e. where LexA concentration is highest. This can be estimated as follows. Previous studies have reported that it takes a single TF molecule on the order of 3 − 5 minutes to find a given binding site on the genome [43, 44]. The number of LexA molecules per cell (in absence of DNA damage) has been estimated as 1100 − 1300 [13, 45], resulting in 550 − 650 dimers [23]. Therefore, when LexA is at its steady-state level, the time-scale for a single unbound binding site to become rebound is predicted to be less than a second. This suggests that in the absence of DNA damage, unbound promoters are rebound so quickly by LexA that it is unlikely that multiple transcription initiation events can occur in between. That is, these previous measurements suggest that production bursts require LexA concentration to decrease substantially.

This raises the question as to what is known about LexA concentration dynamics. It is known that, upon DNA damage events that produce single-stranded DNA, RecA forms filaments on the single-stranded DNA and LexA auto-cleaves itself on these filaments, leading to a decrease in LexA concentration [46, 47]. The decay rate of LexA has been extensively studied under UV-induced damage. However, it is little studied for continuous antibiotics exposure [48]. Moreover, it is challenging to interpret the LexA decay observed in bulk population studies, since this likely represents a heterogeneous mixture of LexA decay and recovery across single cells. Notably, *in vitro* studies reported decay rates on the order of minutes [49–51], and these results agree with several *in vivo* studies that also found LexA concentrations decay within minutes [13, 52]. Most relevant for our current study, a recent single-cell study in cells subjected to low levels of Cipro (i.e. 0.5 ng/mL) found exponential decay of LexA concentration with a half-life of less than a minute [48]. The available data thus suggests that, upon formation of RecA filaments, LexA levels decay 10-100 fold within a few minutes.

These considerations suggest the following picture of the concentration dynamics within single cells (Fig. 6A). Whenever a damage event occurs that leads to a RecA filament, the LexA concentration will start decaying exponentially at a rate *α* (which is larger than 1 per minute, see Table 1). After some period of time the DNA damage will be repaired, RecA filaments will disappear, and the concentration of LexA will recover at a rate *β* (which, again to reduce the number of free parameters, we assume to equal *α*).

**Fig 6.**
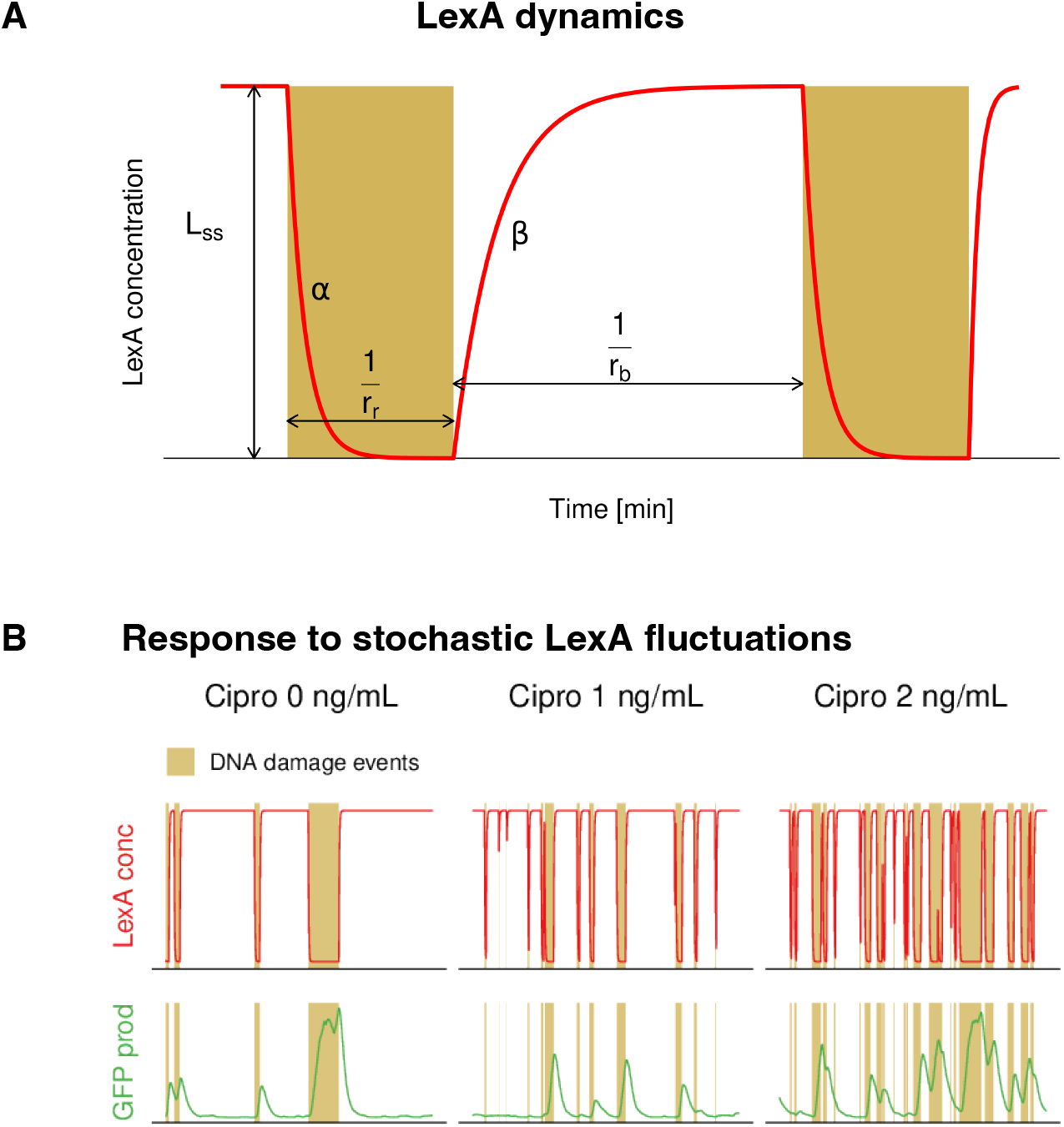
Dynamic model for the induction of the LexA regulatory network. **A**: LexA dynamics in the presence of DNA damage events that leads to the formation of RecA filaments. Without damage (white areas), LexA has a steady state concentration with binding rate *L*_*SS*_. Upon a DNA damage event, LexA concentration starts decaying at rate *α* (yellow areas) until the damage is repaired. We assume damage is repaired at a constant rate *r*_*r*_, and that upon repair, the LexA concentration recovers at a rate *β*. Finally, we assume the DNA damage events occur at a rate *r*_*b*_, which increases with Cipro concentration. **B**: Example simulation results of the dynamic model, where the LexA concentration dynamics of panel A is explicitly taken into account (top row). Production bursts occur when LexA unbinds from a promoter during a DNA damage period when the LexA concentration is low (bottom row). We refer to the Suppl. Mat. section 5 for a detailed description of the simulation algorithm.

We of course do not know the distribution of repair times and this likely depends on the type and extent of the DNA damage. For our model of the damage resulting from relatively low levels of Cipro, we will assume that damage is repaired at a constant rate *r*_*r*_, leading to an exponential distribution of repair times, which is arguably the simplest assumption that can be made for the distribution of repair times. As we will see below, this simple assumption results in a exponential distribution of peak heights, in agreement with our experimental observations.

Together, these considerations define a dynamic non-equilibrium model of the stochastic gene expression dynamics (detailed in Suppl. Mat. section 5) with parameters that are listed in Table 1. Figure 6B shows a representative example of the DNA damage events, LexA concentration, and gene expression dynamics of a LexA target promoter from a simulation of this dynamic model. As already mentioned above, most of the kinetic parameters of this model are known (at least approximately) from previous work (green colored parameters in Table 1), leaving only three unknown parameters: the rate of DNA damage repair *r*_*r*_, the volumic rate *r*_*m*_ at which a given target promoter transcribes when not bound by LexA, and the frequency of DNA damage events at each Cipro concentration *r*_*b*_.

We next investigated whether these 3 remaining parameters of our simple model can be set so as to reproduce the observed burst durations, the exponential distributions of burst heights, and the burst frequencies. We find that this is indeed the case, but that this requires that the repair rate *r*_*r*_ lies in a narrow range. In particular, given that mRNA decay times are typically in the range 2 − 6 minutes and that GFP maturation time is 6 − 8 minutes, even an instantaneous burst of mRNA production results in a peak of volumic production *q* with a duration only a bit less than the observed average peak duration. Simulations of our dynamic model confirm that, to reproduce the observed peak durations, the DNA damage events have to be short-lived on average and the repair rate must lie in the range 0.5 ≤ *r*_*r*_ ≤ 1.0 per minute (Suppl. Figs. S11, S12, and Suppl. Mat. section 6).

Our simulations show that if *and only if* the repair rate lies in this range, the peak heights are also exponentially distributed as observed in our data (Suppl. Fig. S12). In Suppl. Mat. section 6 we provide analytical derivations that explain why peak heights are exponentially distributed in this regime, which we briefly summarize here. Outside of DNA damage events promoters will be almost always bound by LexA. Depending on the strength of the LexA site(s), LexA will stochastically unbind from the promoter a few times each minute, but the rate of rebinding of LexA to its site(s) will be so much higher than the rate of transcription that no burst in mRNA production can occur.

Thus, a transcription burst only occurs whenever LexA unbinds from the promoter during a time interval when the LexA concentration is so low that the rate of LexA rebinding is considerably smaller than the rate of transcription. Because DNA damage is repaired quickly on average, damage events that last long enough for LexA to decay sufficiently may be relatively rare. However, in the subset of damage events that last long enough such that LexA has decayed sufficiently and, in addition, LexA has unbound from the promoter, there is now an exponentially distributed additional amount of time before the damage is repaired. Consequently, the number of mRNAs produced during this time interval will be geometrically distributed, and the fact that these time intervals are generally short (because *r*_*r*_ ≥ 0.5 per minute) also ensures that the resulting heights of the peaks in *q* are simply proportional to the number mRNAs produced (Suppl. Mat section 6).

Our model thus not only explains why peak durations are independent of the promoter and Cipro concentration, it also explains why peak heights are exponentially distributed, and controlled by the ratio of the transcription initiation rate *r*_*m*_ and the damage repair rate *r*_*r*_, which is naturally promoter-dependent, but not dependent on Cipro concentrations.

Note that the burst frequency of each promoter will depend on how often DNA damage lasts long enough such that LexA rebinding rates fall below the transcription rate of the unbound promoter, and that the LexA repressor unbinds from its site. This thus depends non-trivially both on the volumic transcription rate of the promoter *r*_*m*_ and on the strength of its LexA binding site(s). Thus, while the frequency generally increases with Cipro concentration, it does so in a promoter-dependent manner. Although we have no principled way to derive the rate at which DNA damage events occur as a function of Cipro concentration, we find that it is straightforward to set the rate of DNA damage events *r*_*b*_ such that our dynamic model reproduces the observed distribution of peak frequencies as a function of their cut-off (Suppl. Fig. S13).

Finally, although our simple model ignores several features of the stochastic gene expression dynamics in real cells such as the fluctuations in growth rate and the unequal distribution of cell volume at division, Suppl. Fig. S14 shows that our simple model also reproduces the observed distributions of GFP concentration across cells and Cipro concentrations for the recA promoter with reasonable accuracy.

### Promoter response functions as a function of LexA binding site strength and DNA damage durations

In the last sections we have seen that all our experimental observations can be explained by a non-equilibrium model in which production bursts occur when DNA damage events cause short-lived dips in LexA concentration, which occur on time scales that are similar to the time scale at which LexA unbinds from its sites. In this model, the response of different LexA target promoters to DNA damage events will depend in a non-trivial manner on the strengths of the LexA target sites in the promoters, their volumic production rates when unbound, as well as on the number and durations of the DNA damage events. Here we use simulations of our model to explore how the ‘promoter response function’ depends on these parameters.

We first investigated how the promoter output resulting from a single DNA damage event depends on the duration of the event and the strength of the LexA site in the promoter. We set all parameters of our model as in Table 1 and set the transcription initiation rate to *r*_*m*_ = 5 min^−1^*µm*^−1^, which approximately corresponds to the inferred *r*_*m*_ for the RecA promoter. We then performed simulations of the dynamic model under many repetitions of a single DNA damage events of different durations (see Fig. 7A), and measured the distributions of the total number of mRNA molecules produced during the damage event as a function of both the duration of the damage event and the strength of LexA binding sites, as quantified by the promoter’s unbinding rate *L*_off_. Figure 7B shows box-whisker plots of the distribution of promoter output for different *L*_off_ and damage event durations, scaled relative to the total output that would be observed if the promoter had been unbound throughout the entire DNA damage event.

**Fig 7.**
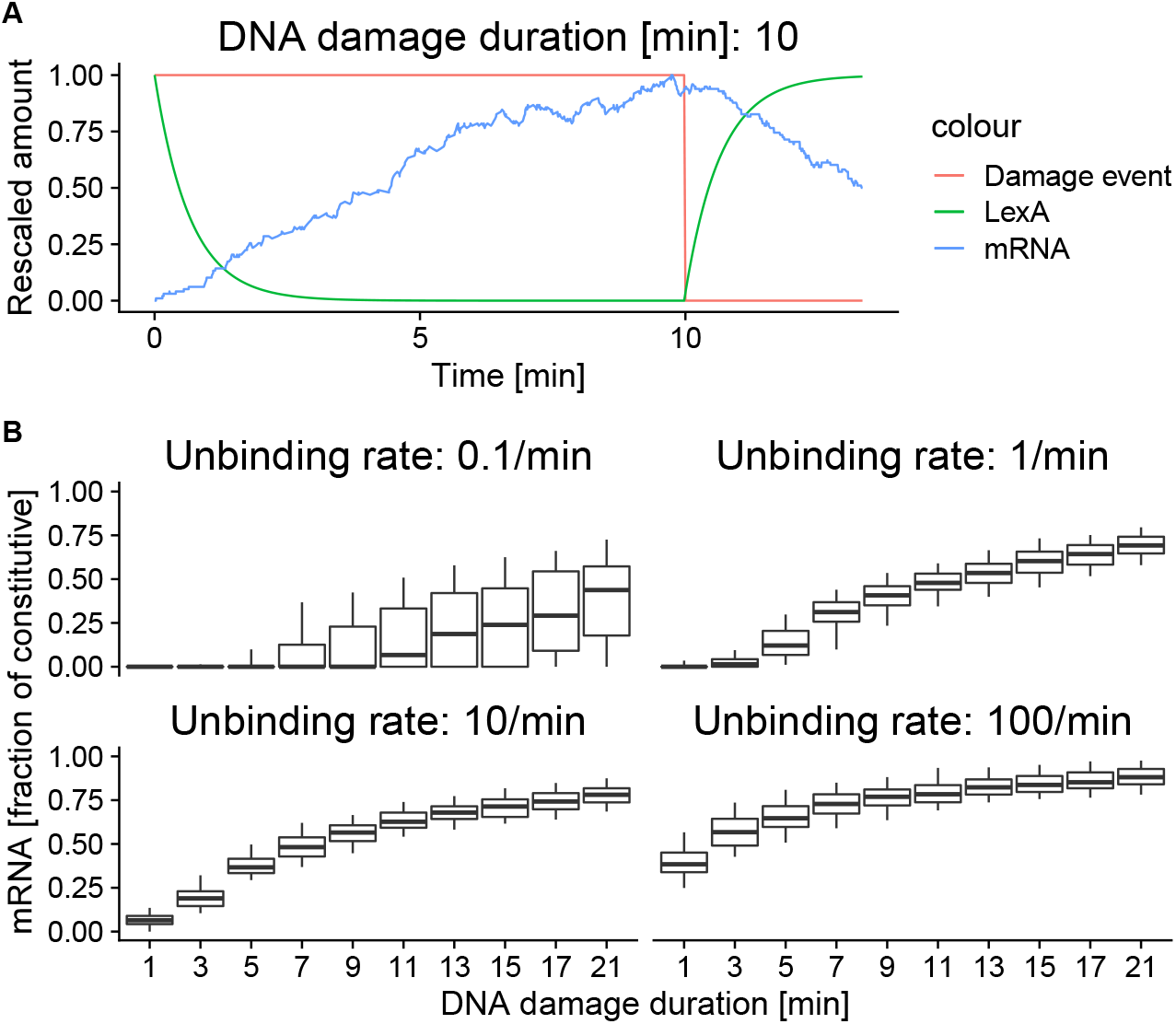
Response of a LexA target promoter to single DNA damage events of different durations. **A**: Example dynamics of DNA damage (red), LexA concentration (green) and mRNA level (blue) for a DNA damage event that lasts 10 minutes. All quantities are rescaled between 0 and 1 for ease of visualization. **B**: Distributions of the amount of mRNA produced (as fraction of the mRNA produced by the fully unbound promoter) as a function of the duration of the DNA damage event (horizontal axis) for LexA target promoters that have different rates of LexA unbinding from 0.1 to 100 per minute (panels). The box-whisker symbols show the 5th, 25th, 50th, 75th, and 95th percentile of the amount of mRNA produced across replicate simulations of DNA damage events of the same duration. We see that unbinding rates in the range 1 − 10 per minute, which correspond to those observed for native LexA sites, give the most sensitive and reproducible response for damage events that last 1 − 20 minutes.

The strongest possible LexA binding site has been reported to have an unbinding rate of 0.1 per minute [18] and we see that, because it takes on the order of 10 minutes for LexA to unbind, promoters with such a LexA site only respond to damage events that last 10 minutes or more (Fig. 7B, top left panel). In addition, we see that for damage events that last 10 − 20 minutes, the response is highly variable. That is, for promoters with such a strong LexA site, the gene expression response is very noisy.

Many native LexA target promoters have binding sites with unbinding rates in the range of 1 − 10 per minute. For promoters with binding sites of these strengths the promoter output increases monotonically, and with less variability, over the entire range of damage durations from 1 to 20 minutes (Fig. 7B, top right and bottom left panels). We see that, the weaker the binding site, the more sensitive the promoter becomes to short damage events. However, for weaker sites the output also saturates more quickly as a function of the damage duration. Thus, the strength of a promoter’s LexA binding site determines over which range of DNA damage durations the promoter goes from not responding to a saturating response. In addition, we also see that the stronger the LexA site, the more variable the responses are across individual damage events. Finally, when the binding site becomes very weak, the promoter starts to become active even in the absence of any damage events, and the range of the response to DNA damage becomes strongly attenuated (Fig. 7B, bottom right panel).

The results in Fig. 7 show that, as expected, the promoter output increases with the duration of the DNA damage. Note that, if the promoter response could be described by an equilibrium model, then the output from the promoter should only depend on the total amount of time that LexA concentration is low. Therefore, the non-equilibrium behavior of the LexA regulon can be illustrated by comparing promoter responses to DNA damage events that have the same total duration, but that vary from one long event to multiple short events. We performed simulations of the dynamic model for a period of 35 minutes in which *n* equally spaced DNA damage events occurred that together have a total duration of 17.5 minutes (Fig. 8A) and again measured the distribution of the total number of mRNA molecules produced as a function of the number of events *n*, and the strength of the LexA binding site (Fig. 8B).

**Fig 8.**
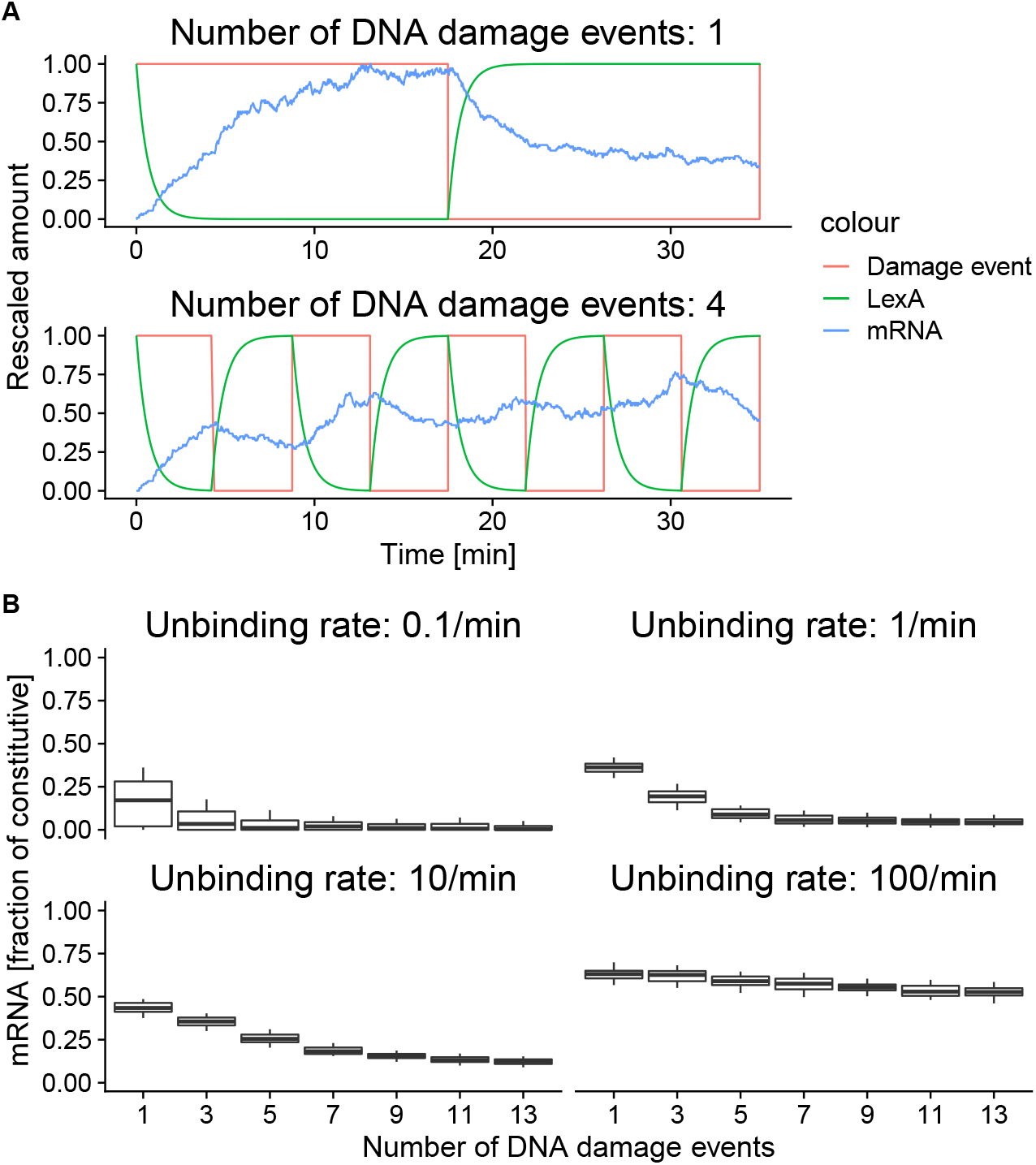
Response of LexA target promoters to different numbers of DNA damage events with the same total duration. **A**: Example dynamics of DNA damage (red), LexA concentration (green) and mRNA number (blue) for either a single damage event lasting 17.5 minutes (top), or 4 damage events lasting 17.5*/*4 = 4.375 minutes each. The quantities are rescaled between 0 and 1 for representation purposes. **B**: Total amount of mRNA produced (as fraction of the mRNA produced by the fully unbound promoter) as a function of the number of damage events (horizontal axes) for different LexA unbinding rates ranging from 0.1 − 100 per minute (different panels). The box-whisker symbols shows the 5th, 25th, 50 (line), 75, and 95th percentile of the mRNA produced across replicate simulations. Note that, since the sum of the duration of the DNA damage events is kept constant, the dependence on the number of events shows that the output from the promoter depends non-linearly on the duration of DNA damage events.

The results show that the total output does not only depend on the total duration of the DNA damage, but is highly dependent on whether the damage is spread over multiple short or fewer longer events, confirming the intrinsically non-equilibrium nature of the promoter response. We find that the promoter response generally decreases with the number of events. For the promoter with the strongest LexA site, there is only a (highly variable) response for the longest events (Fig. 8B, top left panel). For unbinding rates in the regime of the sites in many LexA target promoters, i.e. from 1 − 10 per minute, the response decreases systematically with the number of damage events and disappears when the events become short relative to the average time for LexA to unbind (Fig. 8B, top right and bottom left panels). That is, the unbinding rate of the LexA site sets a lower bound on the duration of the DNA damage events that are detected by the promoter.

The results also illustrate that the dynamic range in promoter response as the DNA damage varies from *n* = 1 event of 17.5 minutes, to *n* = 13 events of 1.3 minutes each, depends strongly on the unbinding rate of the promoter. In particular, the dynamic range is largest for the promoter with unbinding rate 10 per minute because it is able to detected even the shortest damage events, while still not being saturated by the longest event. This illustrates that the unbinding rate of the promoter sets the range of damage event durations over which its response varies. Finally, when the unbinding rate becomes so high that it is on the order of the LexA binding rate, the variation in response becomes highly attenuated because the promoter is active even in the absence of DNA damage (8B, bottom right panel). This illustrates that the LexA concentration in the absence of DNA damage sets a lower bound on the durations of damage events that can be detected.

In summary, the output from a LexA target promoter depends in a non-linear manner on the strength of its LexA binding site(s) and the durations and numbers of DNA damage events. Consequently, the expression levels of different LexA target promoters effectively ‘read out’ different statistical properties of the DNA damage events in the recent history of the cell.

## Discussion

Due to unavoidable thermal noise, gene expression is inherently stochastic at the single-cell level [5] and also the effects of TFs on the expression of their target genes are inherently stochastic, leading to propagation of noise from regulators to their targets [4]. Indeed, recent genome-wide studies have shown that, in *E. coli*, noise propagation drives highly condition-dependent patterns of gene expression noise [6, 7]. However, there is currently little quantitative understanding of how the actions of TFs determine expression responses of different target promoters in single cells.

To study how the induction of a single TF affects the expression dynamics of its target genes in single cells, we here made use of recent technical advances that use fluorescence time-lapse microscopy in combination with microfluidics to allow tracking of cell growth and gene expression in many lineages of single cells with high resolution and quantitative accuracy [26, 27]. Furthermore, we chose the LexA regulon in *E. coli* as object of our study because it combines several attractive features that facilitate the development of realistic quantitative models of its stochastic single-cell dynamics, i.e. 1. it can easily be induced to different levels in the laboratory using low level antibiotics, 2. it has several different target promoters that are all known to be only regulated by LexA, and 3. much is already known about the induction mechanism of the LexA regulon as well as many of its kinetic parameters. In particular, the LexA regulon responds to DNA damage and we monitored the single-cell responses of a collection of target promoters to mild DNA damage using different levels of the antibiotic Ciprofloxacin.

We found that LexA target promoters express in bursts that exhibit very particular quantitative characteristics. While the frequencies of expression bursts systematically increase with Cipro level, their heights and their durations are independent of Cipro level. In addition, the heights of expression peaks are exponentially distributed for all promoters and peak durations have narrow distributions that are identical for all promoters.

We showed that these observations are explained by a simple model in which stochastic DNA damage events cause short transient dips in the concentration of LexA and that bursts occur when LexA unbinds from target promoters during these concentration dips. Crucially, because unbinding of LexA from its target sites (and its subsequent rebinding) occurs on the same timescale as the duration of the LexA concentration dips, the output from each promoter is a complex function of the strength of its lexA binding site(s), its transcription rates when the promoter is either bound or unbound by LexA, and the statistics of the durations of DNA damage events (Figs. 7 and 8).

This is in contrast to popular ‘equilibrium’ models which assume that TF binding and unbinding is sufficiently fast so that the transcriptional output from a promoter can be calculated by averaging the transcription rate over all possible TF binding configurations, each weighted with their thermodynamic equilibrium probability of occurrence [8–10]. Notably, in such equilibrium models the output of a target promoter is a relatively simple logistic function of TF concentration that is characterized by the maximum fold-change and the TF concentration at half-maximum. Consequently, under an equilibrium model there is relatively little variation in the possible responses that different target promoters can give to a change in the concentration of the TF that regulates them. In contrast, the ‘non-equilibrium’ behavior of the LexA regulon causes different target promoters to effectively read out different statistical properties of the DNA damage events in the recent history of the cell.

In this regard it is important to note that our model makes various simplifications. For example, the model assumes that each LexA target promoter can only take on two states, i.e. bound and unbound, and that it only transcribes in the unbound state. However, many native LexA target promoters in *E. coli* have multiple LexA binding sites, so that there are multiple binding states that may all have different rates of transcription. Therefore, in reality the response of each promoter is an even more complex function of the durations and numbers of DNA damage events than in our simple model.

Our data suggest that, in our conditions with only low levels of DNA damaging agent, there are only isolated DNA damage events and LexA levels recover in between. In contrast, when damage events become more frequent or multiple events occur at once, as may occur under UV radiation, the depletion of LexA protein will start feeding back positively on itself and the concentration dynamics of LexA will become much more complex. Although modeling such scenarios are beyond the scope of this work, we believe that the bursting behavior that we observe here, and that is explained by our dynamic model, may also help explain previous observations that under UV damage different LexA target promoters appear to respond at different times [14, 18, 19, 62–64]. This is an important avenue for future study.

Finally, given that the complex promoter response function is determined by the configuration and strengths of LexA binding sites, natural selection may have shaped the response characteristics of different LexA target promoters so as to fine-tune how much induction each promoter should exhibit in response to different types of DNA damage. This raises the question as to whether there are any evolutionary trade-offs in how many different distinguishable responses to the same TF induction can be realized by varying promoter architecture. Our results show that the strength of the LexA binding site sets a lower bound on the length of DNA damage events that can be detected, because LexA is unlikely to unbind during events that are short relative to the unbinding rate. Consequently, weaker sites are much more sensitive to short events. However, this sensitivity requires that, in the absence of DNA damage, the promoters with weak sites are still kept sufficiently repressed, and this in turn requires high expression levels of LexA. Thus, cells ultimately ‘pay’ for the sensitivity and grading in promoter responses by requiring high levels of LexA expression. This suggests that the LexA expression level may reflect a trade-off between cost of expression, and the advantage of allowing different promoters to have distinct responses to different types of DNA damage.

## Supplementary Material

### 1 Data analysis

#### 1.1 Number of GFP molecules vs GFP concentration

As cells grow exponentially to approximately double their size over a cell cycle, the total number of GFP molecules in the cell (as estimated from the fluorescence signal) typically grows in proportion to cell size, while GFP concentration remains roughly constant (Suppl. Fig. S1). In our analyses, we thus decided to focus on the dynamics of GFP concentration (GFP molecules divided by estimated cell length) and how this concentration changes due to fluctuations in the amount of GFP produced per unit time and volume of the cell.

**Fig S1.**
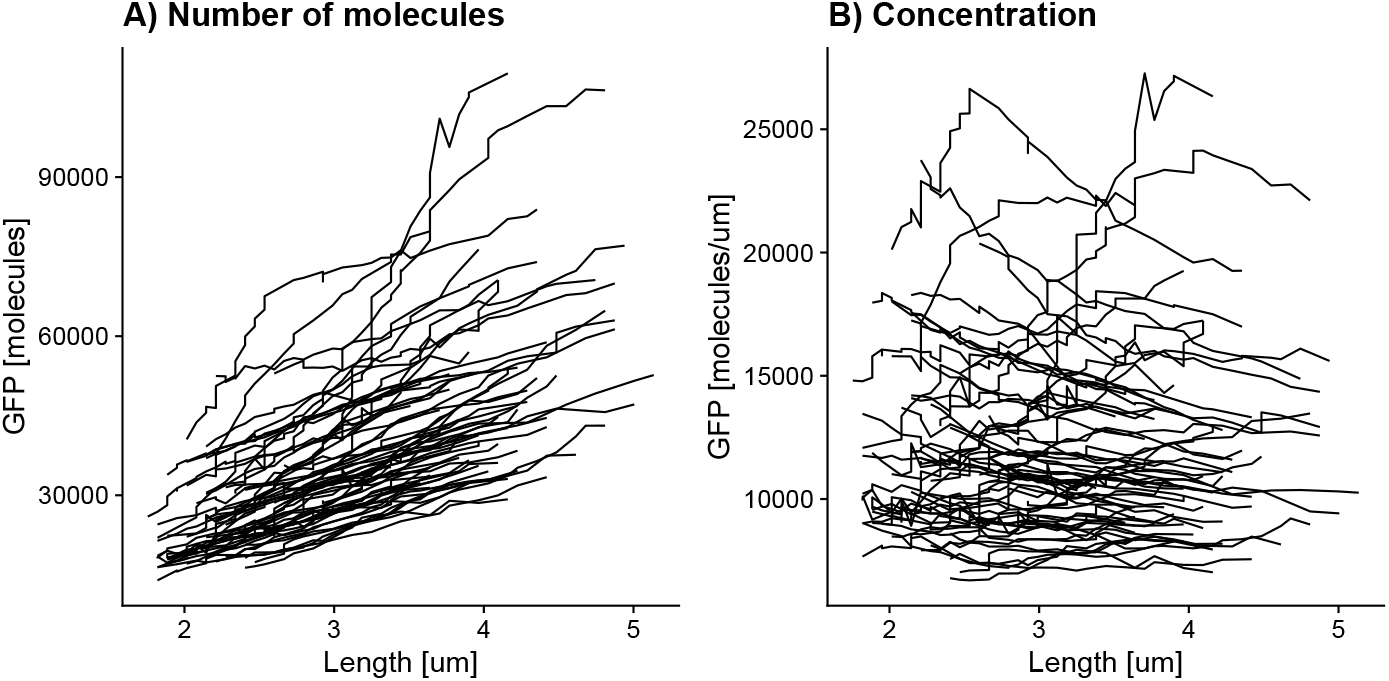
Total GFP molecule numbers are proportional to cell size. **A**: Total GFP number as function of cell length for a selection of cells expressing GFP from the synthetic high promoter. Each line corresponds to one cell cycle. **B**: The same data but now showing GFP concentration (GFP molecule number per unit cell length) versus cell length. Notice there is no systematic correlation between GFP concentration and cell length.

#### 1.2 Day-to-day variation in absolute expression levels

We noticed that there were clearly measurable differences in GFP levels from experiments done on different days, even for the reporters with constitutive synthetic promoters. To illustrate this, Suppl. Fig. S2A we show the average GFP concentration of the Synthetic High promoter across 6 different experiments (horizontal axis) and stratified into 7 different time segments of the experiments (panels). It is clear that there is substantial variation in GFP concentration across experiments, but the fact that the patterns in each panel look similar suggests that, within each experiment, GFP concentrations are relatively constant across the 4 hour time segments.

We normalized GFP concentrations by dividing by the average GFP concentration across all cells and time points within a given experiment, and confirmed that this indeed results in constant expression across time for each experiment (Fig S2B). In particular, the expression variation is now within the estimated error bars across days and time segments. Thus, it appears that subtle variations in the growth conditions on different days lead to measurable changes in absolute GFP levels, but that the relative concentrations across time within an experiment are highly reproducible.

**Fig S2.**
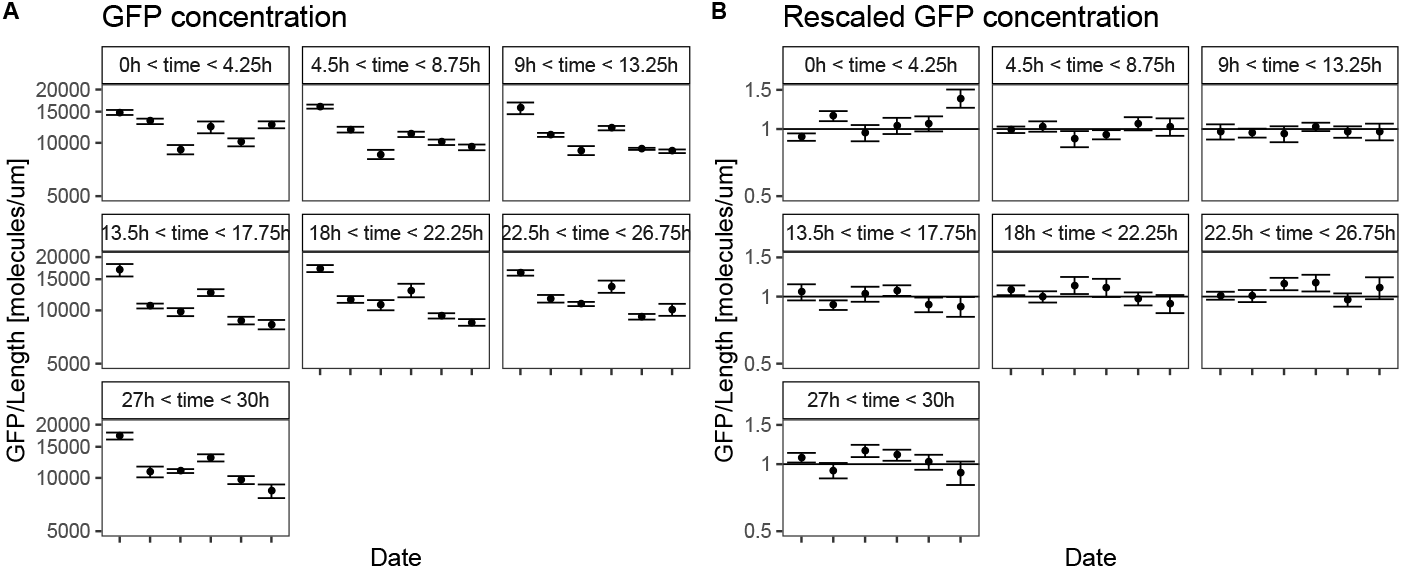
Average GFP concentration. **A**: Average GFP concentration as driven by the constitutive Synthetic High promoter across a population of cells from experiments performed on different days (horizontal axis), and stratified by time bins of 4.5 hours segments of each experiment (one panel per time segment). Note that clear day-to-day variation in average GFP levels is evidendent, and that expression levels are relatively constant across different time segments. **B**: Same as panel A, but we divided each concentration by the average GFP concentration over the entire day. The results show that, after this normalization, expression levels are approximately constant across time on all days.

As a further confirmation, Fig. S10 shows the average GFP concentration in glucose and the fold changes upon treatment with Cipro for all the promoters, and shows that for recA and polB, the normalized concentrations also agree across different measurement days.

#### 1.3 Adaptation to the Mother Machine

Increasing the illumination of the cells allows for increased accuracy and sensitivity in the GFP measurements. On the other hand, when illumination levels become too high, they become toxic to the cells. Indeed, we suspect that the stochastic induction of the recA promoter seen in recent mother machine experiments may be the result of illumination leading to low levels of DNA damage [23, 24]. Compromising between accuracy and phototoxicity we chose an illumination where the cells grow only marginally slower than without illumination.

Measuring the distribution of growth rates as a function of time (obtained by local linear fits of log cell length versus time) we observed that the median growth rate of cells drops during the first two hours of the experiment, presumably due to the cells adapting to the illumination (Suppl. Fig. S3). To exclude effects from this transient adaptation phase, all measurements from cells born in the first two hours were discarded for the analyses presented in this work.

## 2 Inferring the volumic GFP production

Let *G*(*t*) be the amount of GFP inside a cell at a time *t*. We denote by *q*(*t*) the amount of GFP that is created per unit volume and per unit time at time *t*. We assume that GFP fluorescence decays at a rate *β*, which represents the combined effects of bleaching and protein decay. We thus have for the dynamics of *G*(*t*):

**Fig S3.**
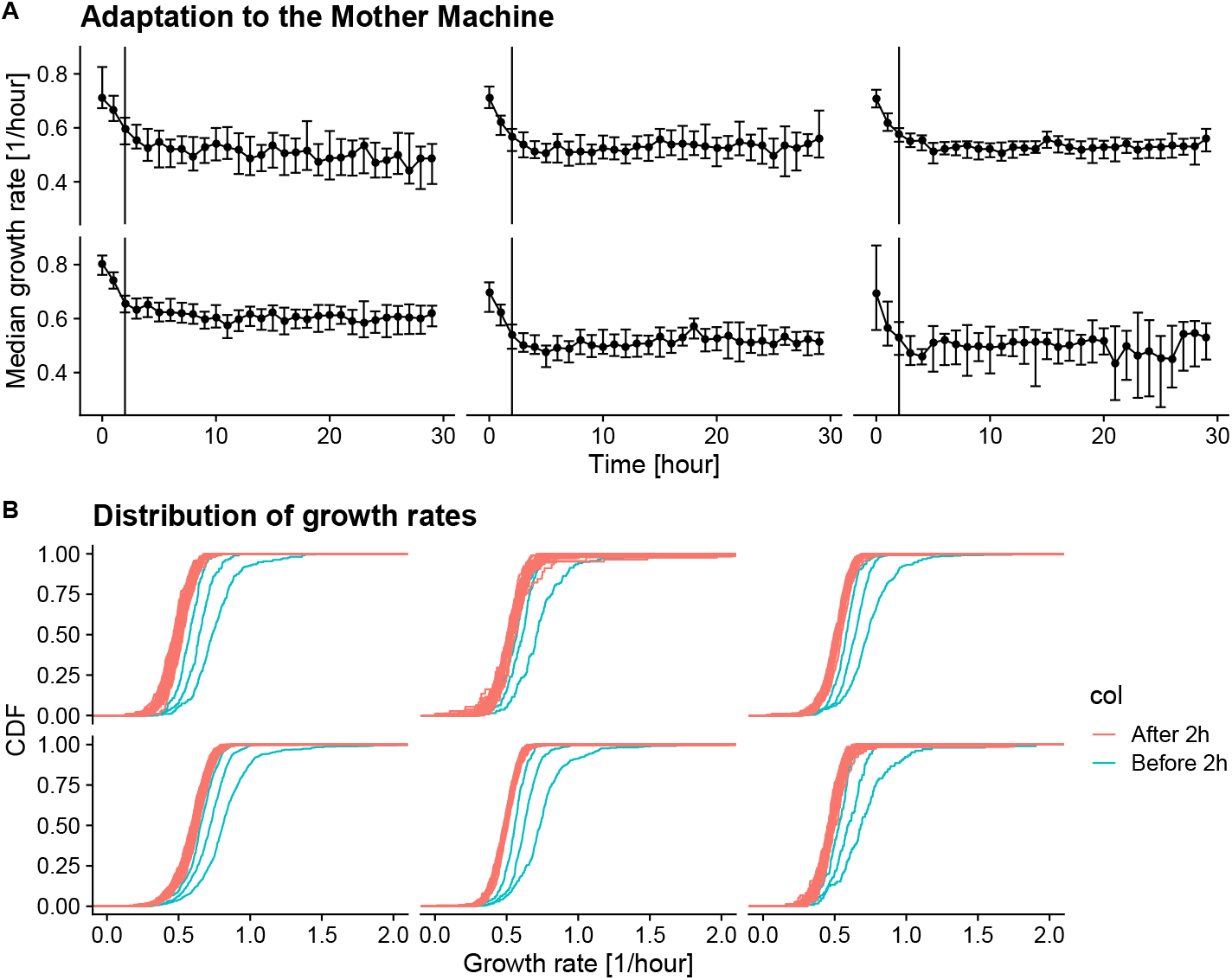
Cells adapt to illumination during the first two hours. **A**: Median growth rate of the cells during different periods of the experiment (horizontal axis) as measured on different days (different panels). After the illumination commences at time 0 hours, the growth rate first decreases and then stabilizes after approximately 2 hours. Error-bars correspond to two standard-deviations of the posterior of the median. **B**: Cumulative distributions of single-cell growth rates stratified by time bin. The blue lines show the distributions coming from time bins earlier than 2 hours and they are clearly higher than the red distributions coming from time bins after 2 hours. Each panel corresponds to a different biological experiment.

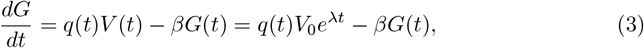

where in the second equation we have assumed that cell volume grows exponentially at a rate *λ* starting from volume *V*_0_ at time *t* = 0. We can solve this equation by rewriting it as

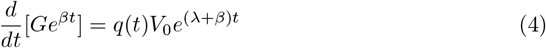

Integrating and setting the initial condition *G*(*t*_0_) = *G*_0_ we find

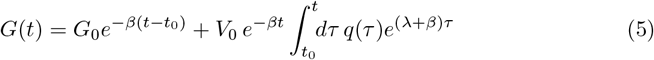

In order to estimate a volumic production rate *q*_*i*_ at each measurement time point *t*_*i*_ we will make the approximation that *q*(*t*) is approximately constant over a time window of *T* = 5 time points centered on *t*_*i*_ and estimate this *q*_*i*_ from the measurements of total fluorescence in this time window.

If we set the volumic production rate *q*(*t*) to a constant *q* for a short time window, we find for the total GFP content during this time window:

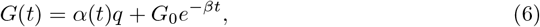

where we have defined

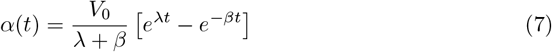

Let 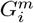 correspond to the measured total GFP fluorescence at each time point *t*_*i*_ in the short time window and assume that each measurement 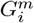 corresponds to the true value *G*_*i*_ plus Gaussian measurement noise of (unknown) standard-deviation *σ*. That is, we assume

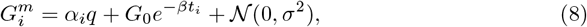

where

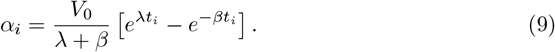

To estimate the *α*_*i*_ for our time window of *T* = 5 time points, we use a slightly larger window of 9 time points centered on the same time point and perform a linear regression of log cell-length versus time to estimate the growth rate *λ* and the cell length *V*_0_ at the start of the time window. We further define 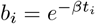, using the total decay plus bleaching rate *β* that we estimated previously [26].

Given the estimated values of the *α*_*i*_ and *b*_*i*_, the probability of the data is given by

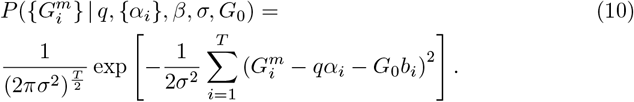

To remove the dependence on the unknown starting GFP level *G*_0_ and the unknown size of measurement errors *σ*, we marginalize over these variables with a uniform prior for *G*_0_ and a scale prior *dσ/σ* for *σ*. Finally, assuming a uniform prior for *q* as well we find that the posterior for *q* is given by the Student-t distribution

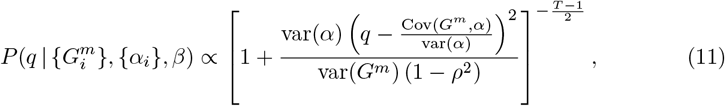

where we defined the variances, covariance, and correlation

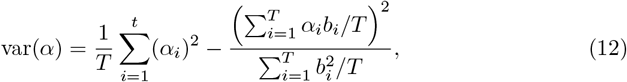

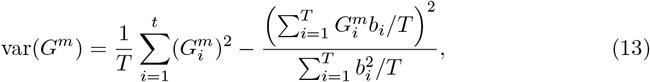

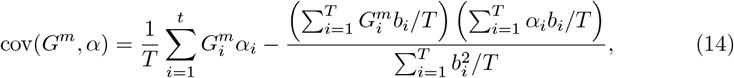

and the correlation coefficient

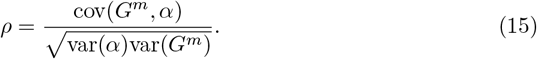

Note that, because the effects of bleaching are incorporated through the dependence on the *b*_*i*_, these definitions of the variances and covariances are slightly different from the usual expressions.

From this we have the maximal posterior estimate for *q*:

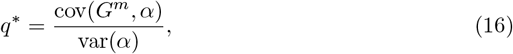

and approximate error-bar

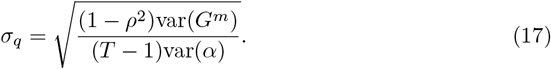

Note that these results are easy to interpret. The best estimate for *q* is simply the usual best estimate for a linear model. The error goes to zero as the correlation *ρ* goes to one, which occurs when the measurements fall on a perfect straight line. The error-bar is further proportional to the standard-deviation of the measured GFP fluorescences *G*^*m*^ and inversely proportional to the standard-deviation of the *α*_*i*_, which means that if the measurements span a larger interval in *α*, the estimate becomes more accurate. Finally, the error depends as usual on the inverse square-root of the number of independent measurements (*T* − 1) (since one independent measurement was lost when marginalizing over *G*_0_).

We thus divide the entire dataset into sliding windows of *T* time points each and for each window estimate a local value of *q*. We checked that reducing the width of the windows below *T* = 5 had no significant effect on the peak statistics, apart from slightly reducing the peaks width.

## 3 Identification of peaks in volumic production

### 3.1 Peak finding method

We observed that for all target promoters that show an induction by LexA (i.e. recA, lexA, recN, polB, and ftsK), the distribution of volumic production rate *q* has an exponentially distributed tail, as shown in Suppl. Fig. S4. Visual inspection of the time traces of *q* confirmed that values out in these tails correspond to bursts in GFP production. In contrast, the distribution of volumic production of the synthetic high promoter does not show such an exponential tail (Suppl. Fig. S4).

**Fig S4.**
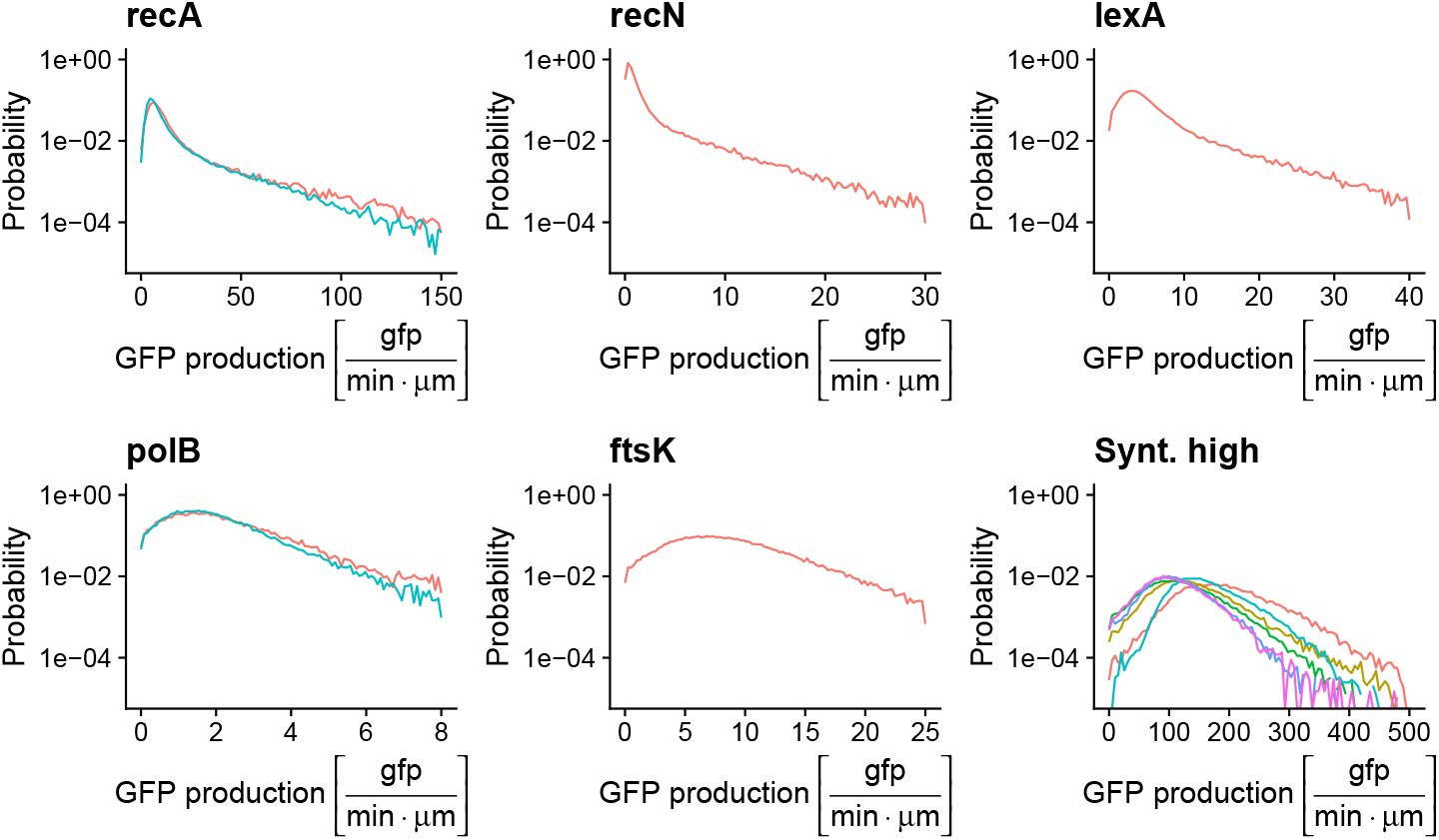
Distribution of volumic GFP production *q* for the LexA target promoters and the Synthetic high promoter. Probability densities of the estimated volumic production rates *q* for each promoter. The *y*-axes are shown on logarithmic scales so that exponential tails appear as straight lines. The production rates *q* were pooled from the entire experiment, regardless of the Cipro concentration. Note that for LexA targets all distributions exhibit an initial peak at low *q* followed by an exponential tail of large *q* values, which inspection suggests corresponds to short-lived peaks in production. In contrast, the synthetic high promoter does not exhibit a similar exponential tail in *q*.

For each LexA target promoter we identified peaks in the time traces of the volumic GFP production rate *q*(*t*) by finding all segments of consecutive time points for which *q*(*t*) is over a cut-off value *q*_*c*_, where *q*_*c*_ is taken to lie in the exponential tail for the corresponding promoter.

We then used the following algorithm:

1. Find all contiguous time segments for which *q*(*t*) *> q*_*c*_.
2. For each such segment, find all local maxima of *q*(*t*) within the segment.
3. For each local maximum, we extract a segment of data points around the maximum to fit it to a Gaussian shaped peak. To meaningfully fit a Gaussian-shaped peak, the segment needs to capture the curvature around the peak. We create a segment by including contiguous data points to the left and right of the local maximum for as long as their *q* values keep decreasing, and as long as the *q* values are at least a fraction *f* = 0.25 of the maximum. For each maximum we thus include a segment of falling *q* values to the left and right, until *q* drops to a quarter of its maximum value.
4. Before we fit the *q* values in the segment to a Gaussian, we first perform two checks: The maximum must not lie at the edge of the segment, and the segment must have at least 3 data points in it. Otherwise the maximum is discarded.
5. For each segment of consecutive *q*(*t*) measurements [*q*(*t*_1_), *q*(*t*_2_), …, *q*(*t*_*n*_)] around a maximum, we fit a Gaussian by minimizing the sum of squared deviations

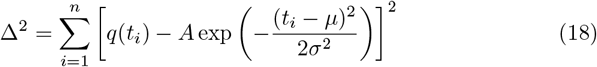

with respect to *A* (height of the peak), *µ* (time of the peak maximum) and *σ* (peak width). In this way we obtain a set of extracted peaks for each Mother Machine experiment, together with a time *µ* of each peak’s occurrence, its height *A*, and its duration *σ*.

### 3.2 Peak height distributions have exponential tails

We next set out to estimate the average peak height *h* of the exponential tail of the distribution for each promoter. In particular, we assume that for peaks with height *A* larger than a cut-off *q*_*c*_, the distribution is exponential, that is

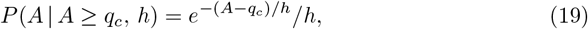

with *h* the average peak height. Imagine that we have *n* peaks with ⟨*A*⟩ ≥ *q*_*c*_ and that 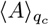 is the average height of these *n* peaks. It is easy to show that the maximum likelihood fit of average peak height *h* to the distribution (19) is given by setting *h* equal to

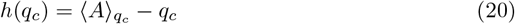

with an error-bar of

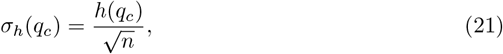

i.e. standard-deviation of the posterior distribution.

To the extent that the tail of the distribution of *q* is really exponential, we should find that the average heights *h*(*q*_*c*_) are relatively insensitive to *q*_*c*_, when the cut-off lies in the exponential tail of the distribution. Supplementary Fig S5 shows that this is indeed the case. Also, the distribution of heights above *q*_*c*_, as shown in Fig 3 of the main text, clearly confirms that the peak heights are exponentially distributed.

**Fig S5.**
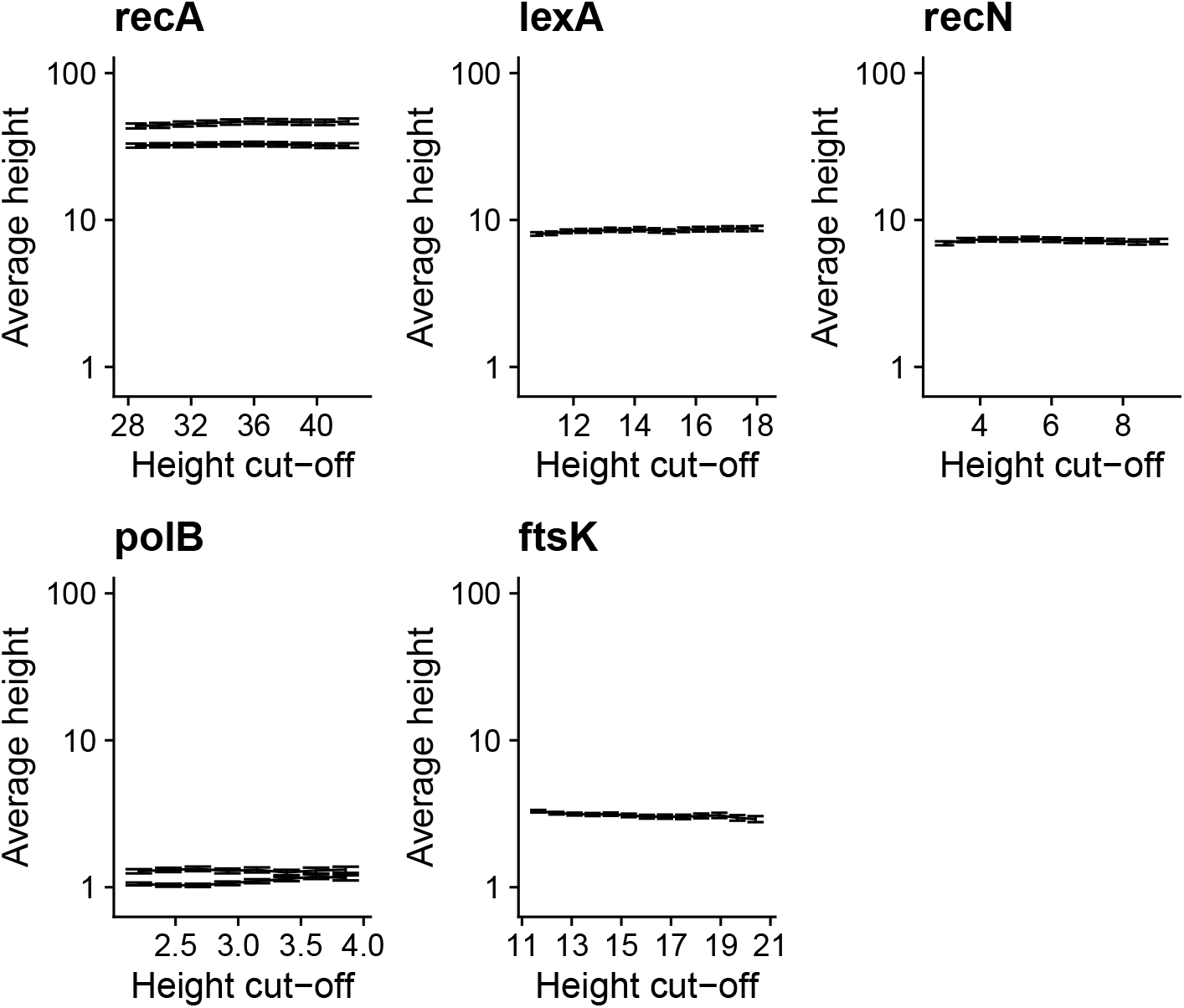
Average peak height *h*(*q*_*c*_) as function of the cut-off *q*_*c*_. Estimated average height *h*(*q*_*c*_) as a function of the cut-off on peak height *q*_*c*_ for each of the target promoters. The error bars show estimated height *h*(*q*_*c*_) plus and minus one standard-deviation *σ*_*h*_(*q*_*c*_). The cut-off *q*_*c*_ for each promoter is chosen to lie in the exponential tail of Suppl. Fig. S4. Note that the estimate of *h* is insensitive to the cut-off *q*_*c*_. For recA and polB estimates from experiments on two separate days are shown.

### 3.3 Peak height and width distributions for polB and ftsK

**Fig S6.**
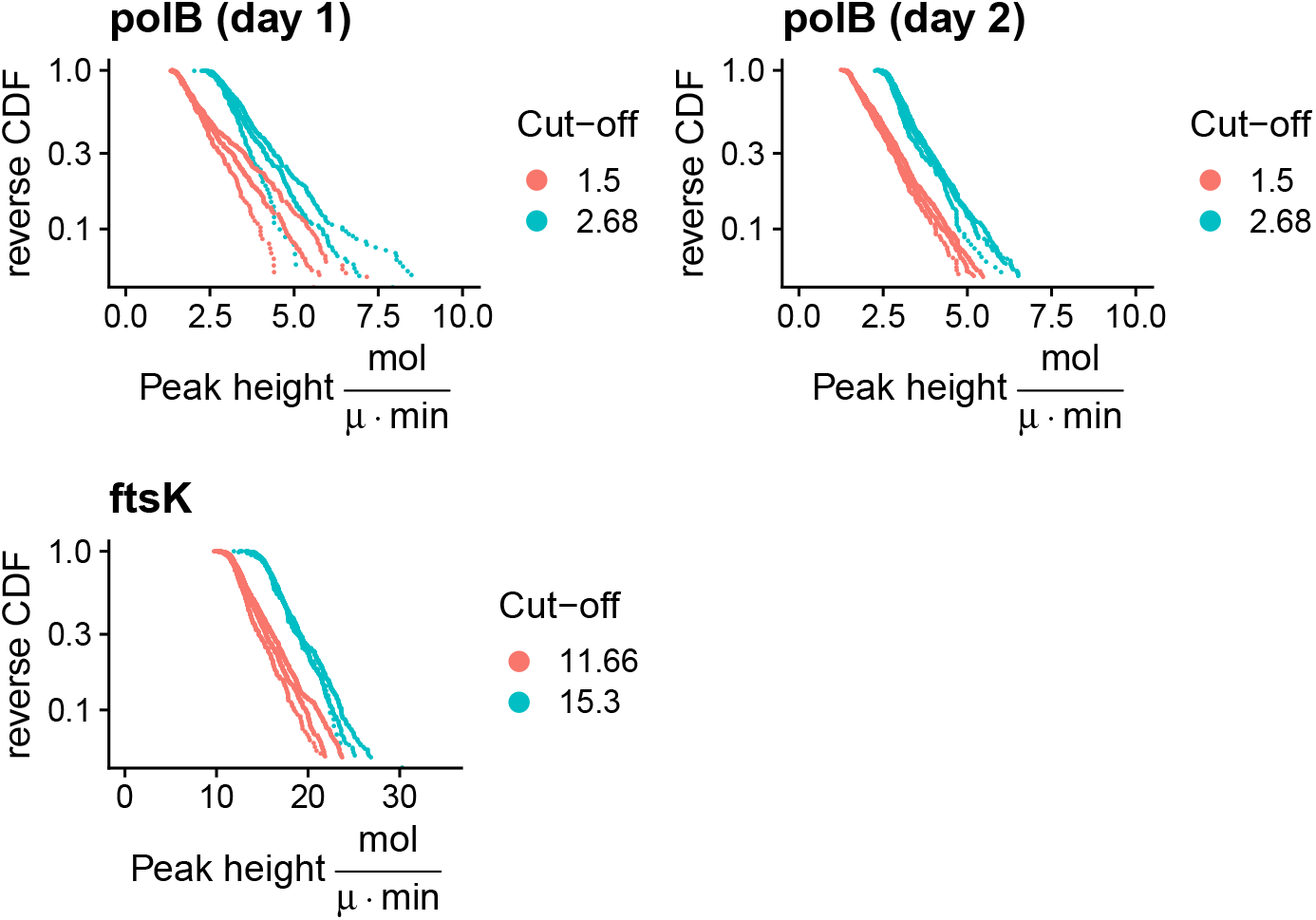
The peak height distribution is exponential and independent of the cutoff *q*_*c*_. The peak heights for polB and ftsK are exponentially distributed for two different values of the cut-off *q*_*c*_ in the range of the exponential tail (Suppl. Fig. S4). Each color represents a different cutoff. Curves of the same color represent the distributions of *q* during the three experimental segments with different antibiotic concentrations. This figure complements Fig. 3 of the main text.

In Suppl. Figs S6 and S7, we show that the distributions of heights and durations of the peaks for the polB and ftsK promoters also exhibit the same features we observed in the main text for recA, recN and lexA. That is, the heights are exponentially distributed and depend only on the promoter, while the durations are independent of both the promoter and the Cipro concentration. The figures also show the distributions for two different values of the cut-off *q*_*c*_ in the range where the GFP production is exponential (see Suppl. Fig. S4).

### 3.4 Peak frequencies increase with antibiotic concentration

As explained in Section 3.1, we identified peaks in the time traces of the estimated volumic GFP production rate *q*(*t*) by finding all segments of consecutive time points for which *q*(*t*) is above a cut-off value *q*_*c*_. The frequencies of the peaks, defined as number of peaks divided by total time, of course depends on the chosen cut-off *q*_*c*_, However, since the heights are exponentially distributed, we expect that the number of identified peaks, and hence the frequencies, are also exponentially decreasing with the cut-off. This behavior is indeed verified in Suppl. Fig. S8, where the cut-off used for each promoter runs over the exponential tail of *q*(*t*) (see Suppl. Fig. S4). In order to define an effective peak frequency that is independent of the cut-off *q*_*c*_, we extrapolate the exponential curve to *q*_*c*_ = 0 for each promoter. Note that, by effectively assuming that peaks can be arbitrarily shallow this effective frequency overestimates the frequency of true expression bursts. The estimate is solely intended to compare frequencies of different promoters and different time segments in a manner that removes the dependency on the cut-off *q*_*c*_.

**Fig S7.**
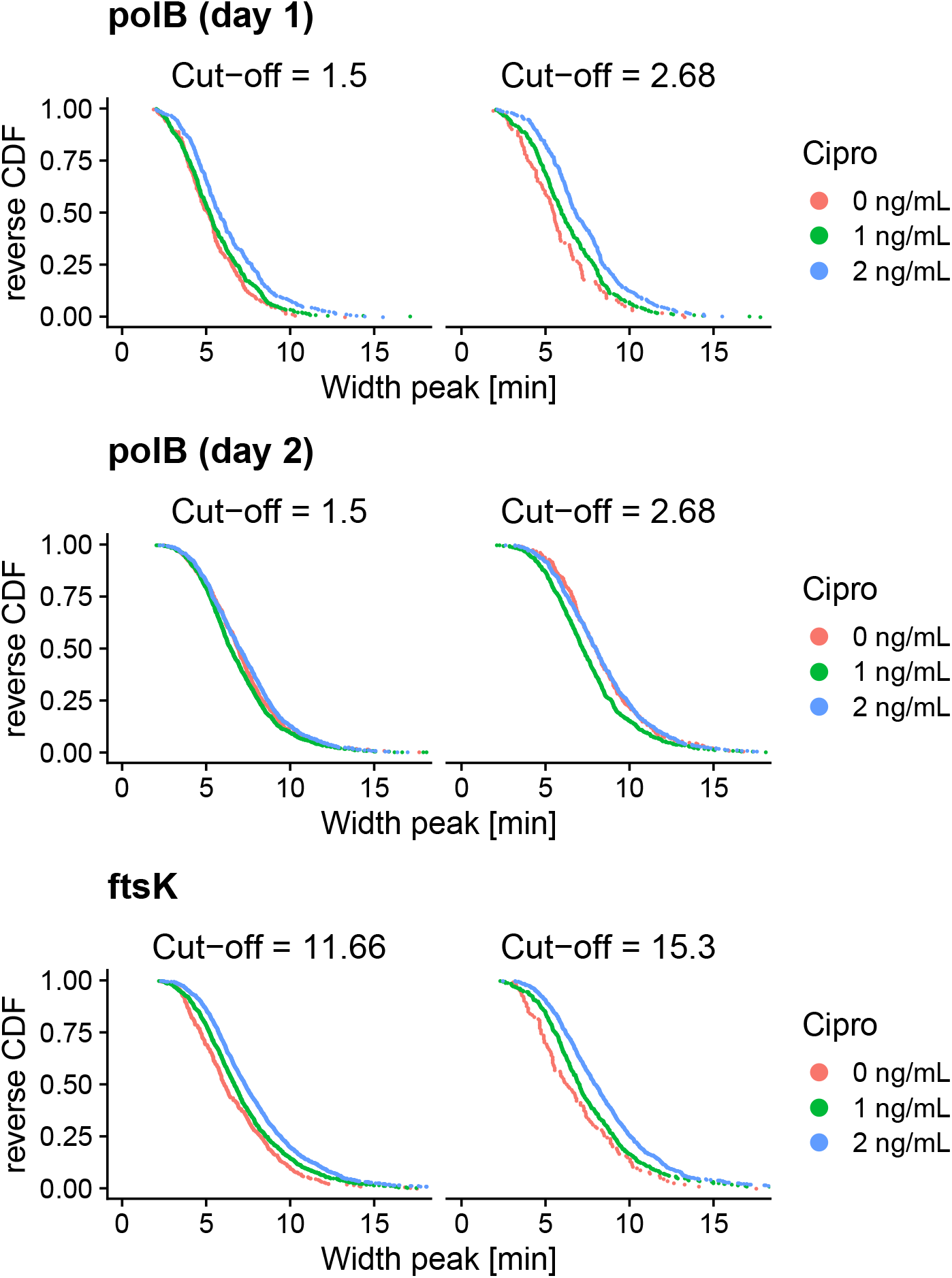
Peak durations do not depend on the promoter, Cipro concentration, or the cutoff *q*_*c*_. Reverse cumulative distributions of the peak durations for polB and ftsK for two different values of the cut-off *q*_*c*_ (different panels) in the range of the exponential tail (Suppl. Fig. S4), separately for each Cipro concentration (different colors). This figure complements Fig. 3 of the main text. Note that for polB results from experiments on two different days are shown.

This extrapolation is done by simply fitting the observed log-frequencies to linear functions of *q*_*c*_ and inferring the intercepts. Notice that for each promoter there are three lines, corresponding to the log-frequencies of the peaks in the three different Cipro concentrations. However, the slope for each Cipro concentration is expected to be the same, since the height distribution is independent of the Cipro concentration. Therefore, we fit each promoter using a maximum likelihood model where the slope is constrained to be the same for the three different concentrations of Cipro. In particular, let’s suppose we have a set of points from *N* linear models indexed by an index *i*, with the same slope *m*, but different intercepts *q*_*i*_. Then

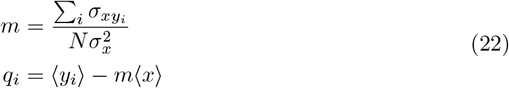

where *i* runs over the *N* lines and we assumed that the independent variables *x* (the Ciprofloxacin concentrations) are the same for all the lines.

The results of the fits are shown in Suppl. Fig. S8, while the inferred peak frequencies are reported in the main text in Fig. 4.

## 4 Effects of Ciprofloxacin on cell growth and gene expression

### 4.1 Single cell growth rates are not affected by Ciprofloxacin

The concentrations of Ciprofloxacin used in our experiments are low enough not to significantly affect the physiology of the cells. Indeed, the distribution of single-cell growth rates is not meaningfully affected by the Ciprofloxacin, as shown in Fig. S9.

### 4.2 Different LexA target promoters exhibit different levels of induction

For each promoter studied in this work, we measured both the average GFP concentration when growing without Ciprofloxacin (Suppl. Fig. S10, top panel) as well as its average fold-change as a function of time along the experiment (Suppl. Fig. S10, bottom panel). Although ruvA, uvrD and umuD are reported to be regulated by LexA, they did not show any induction in their gene expression in our experiments. It is possible that these promoters are only induced at more severe levels of DNA damage, which could cause more prolonged periods of decrease in LexA expression than in our experiments.

## 5 Simulation of gene expression dynamics in growing cells

To rigorously test the predictions of different models for the stochastic gene regulation of the LexA targets we developed a simulation framework that tracks a lineage of cells as they grow, divide, and stochastically express from a LexA target promoter. The algorithm, which we implemented in C++, implements the steps depicted in Fig 5A of the main text:

**Fig S8.**
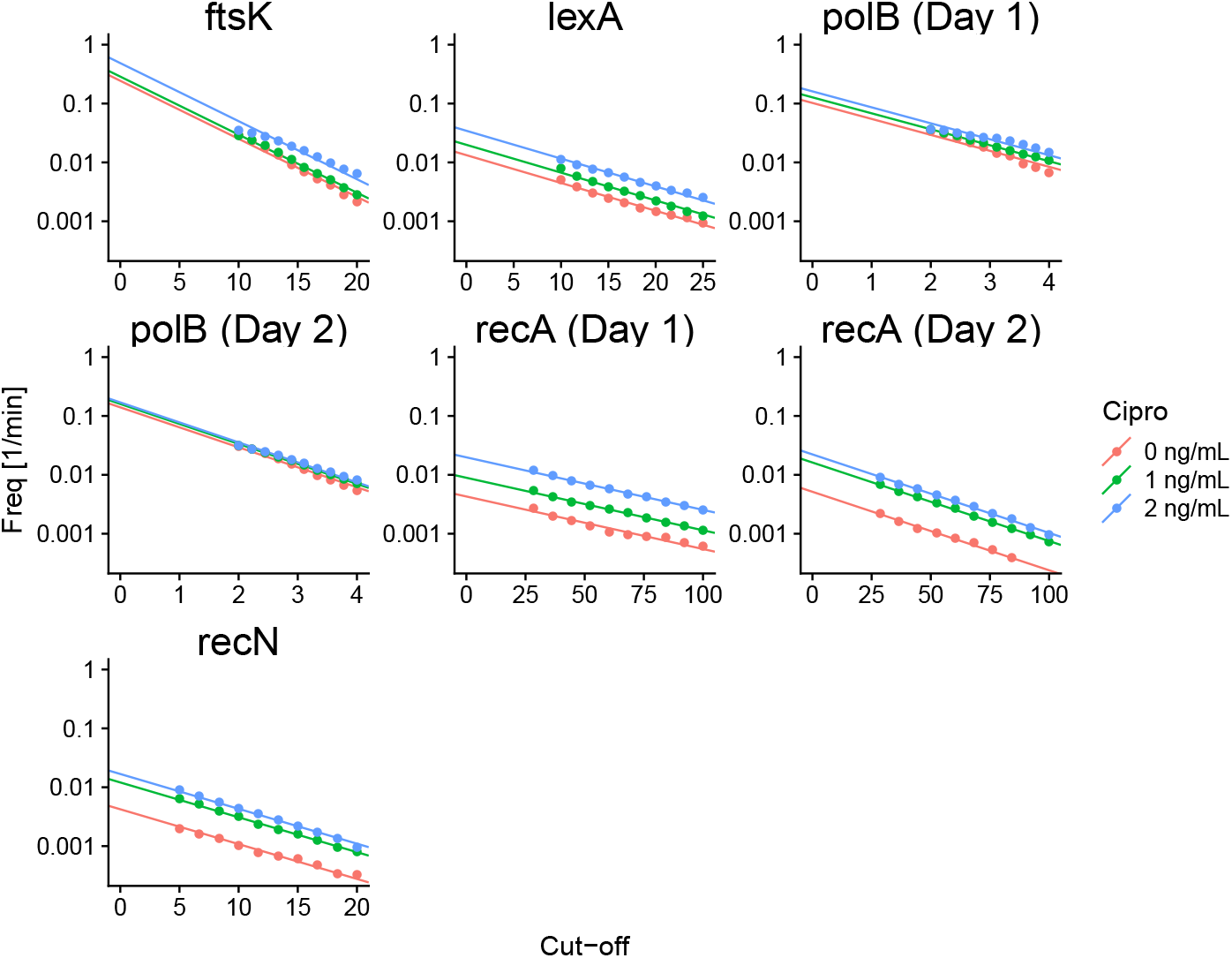
Peak frequency as function of the cutoff *q*_*c*_. Each panel shows the experimentally observed frequencies of production peaks as a function of cut-off *q*_*c*_ for different Cipro concentrations (colors) for a given promoter and replicate experiment (indicated at the top). Peak frequencies decay exponentially with cut-off *q*_*c*_ for each promoter and increase with Cipro concentration. To define an effective peak frequency that is independent of the cut-off *q*_*c*_ we fit exponential functions (solid lines) and extend the curves to *q*_*c*_ = 0. All vertical axes are shown on a logarithmic scale. Note that recA and polB have been measured in duplicate on separate days. Note also that in the fitting we impose that a given promoter has the same exponential dependence on cut-off in all Cipro concentrations, i.e. equal slopes in the log-linear plot.

**Fig S9.**
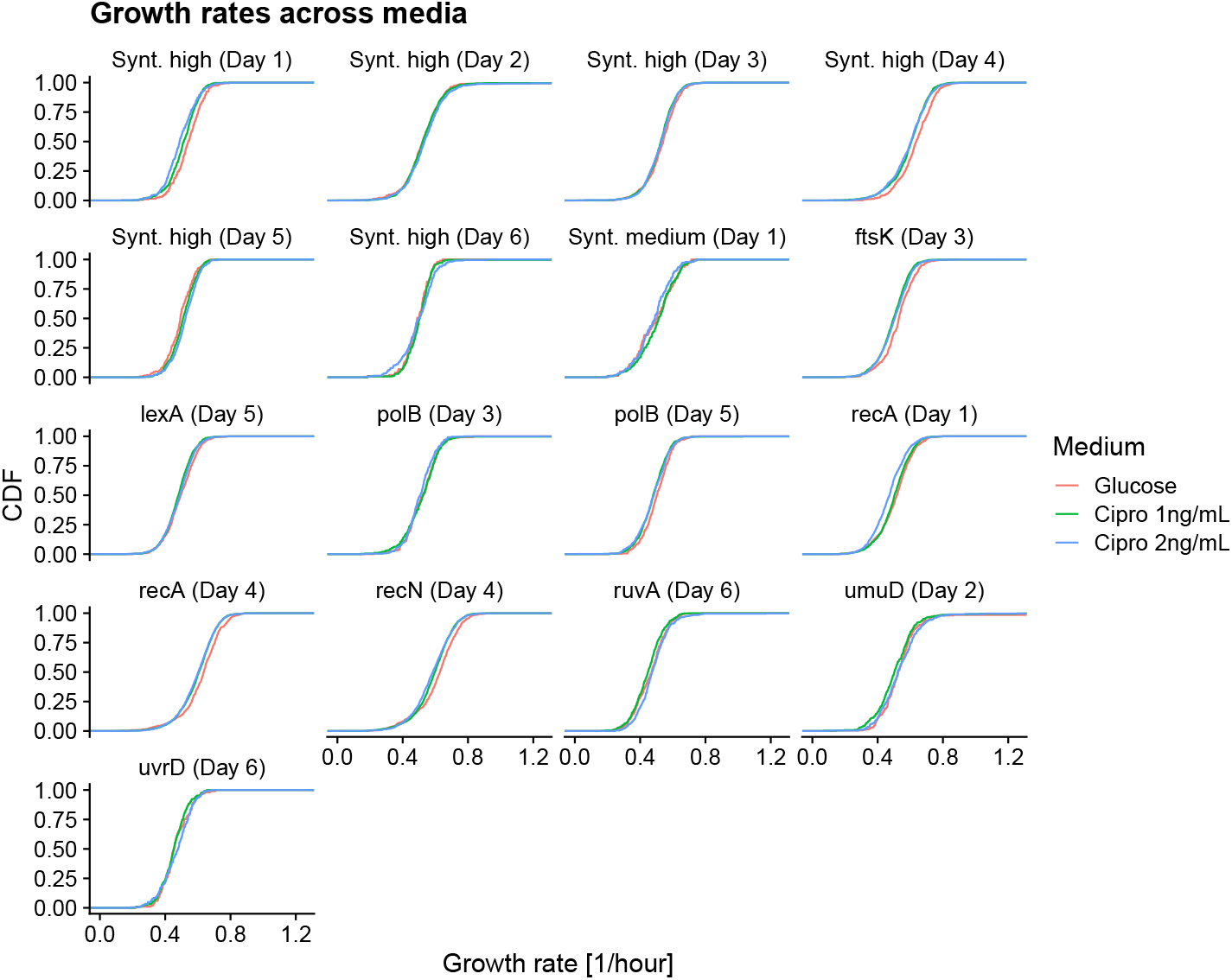
Single-cell growth rates are not affected by Ciprofloxacin. For each observed single cell cycle in each experiment, a growth-rate was estimated by a linear fit of log cell length against time. Each panel shows the cumulative distribution of single-cell growth rates for a given promoter measured on a given day. The different colors represent the three different concentrations of Ciprofloxacin used during each experiment. Note that the growth rates are virtually independent of the Cipro concentration and promoter.

**Fig S10.**
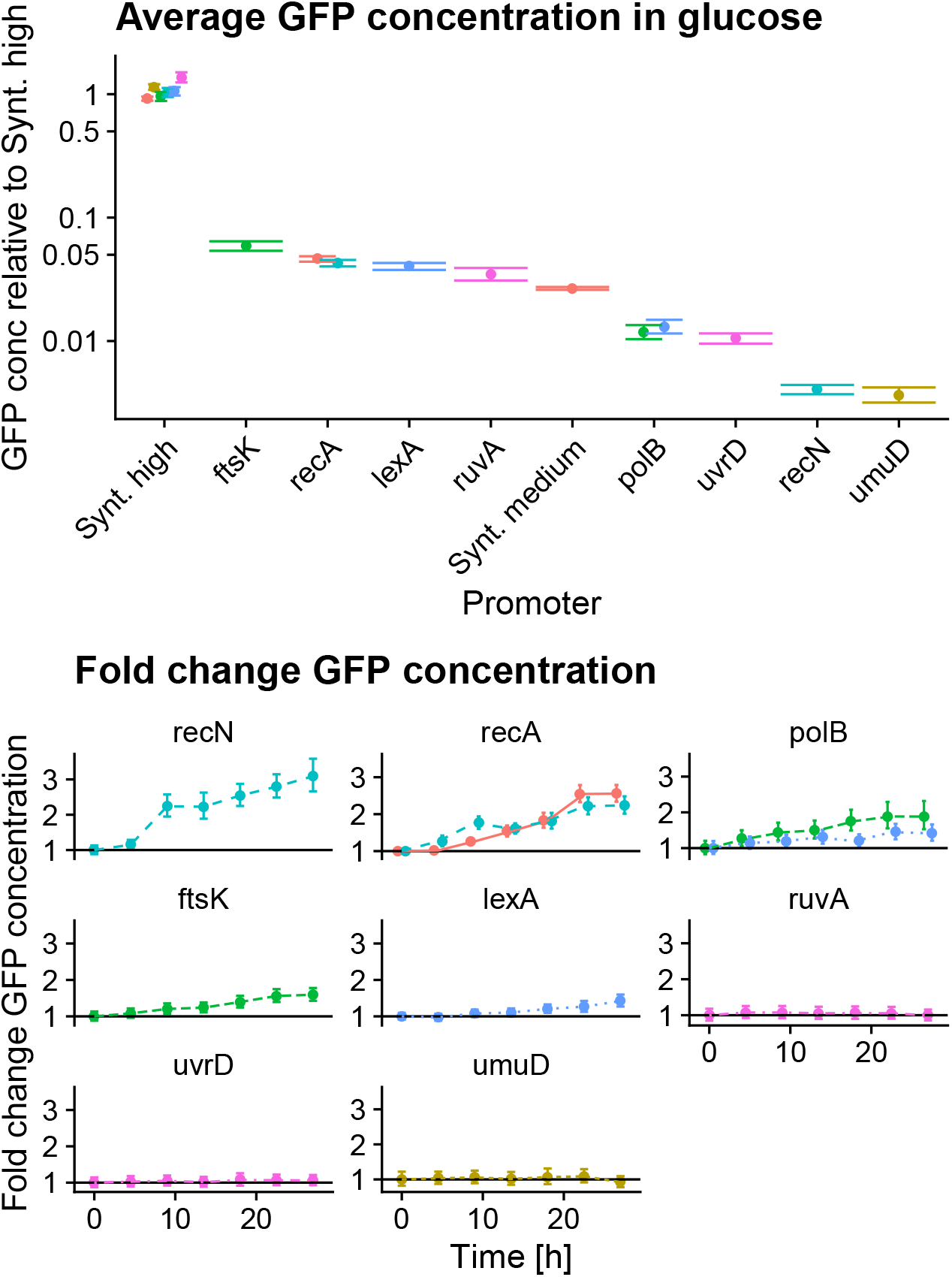
Population average expression levels under treatment with Ciprofloxacin. *Top*: Average GFP concentration for the different promoters studied in this work in conditions without Ciprofloxacin. Note that, in order to take small day-to-day fluctuations in absolute fluorescence levels into account, the basal expression is reported relative to the average GFP concentration of the synthetic high promoter on the same day. Note also that polB and recA have been measured on two different days and different colors represent different experimental dates. Error bars correspond to mean ± one standard-error. The vertical axis is on a logarithmic scale. *Bottom*: Fold change in population average GFP concentration as a function of time during the experiment, with no Cipro for 0 − 6 hours, 1 *ng/mL* for 6 −18 hours, and 2 *ng/mL* for 18 − 30 hours. Error bars correspond to mean ± one standard-error. Note that for ruvA, uvrD, and umuD we did not detect any induction.

1. The simulation is initialized from a very small cell length (2*nm*) whose precise value is not important as the dynamics reaches a stationary cell length distribution relatively quickly, and we discard data from the initial transient phase.

2. Cell cycles follow the adder model [31, 32] and at the beginning of each cell cycle, an added length Δ*L* is drawn from a Gaussian distribution with mean 1.86 and standard deviation 0.35 micrometer, which correspond to the mean and standard-deviation of cell length observed in our experiments. The cell then grows exponentially with a constant growth rate of 0.009/min (matching the median doubling time of 76min we observed in the experiments), until it the added length Δ*L* is reached.

3. After adding a length Δ*L*, the cell divides into two daughters with each exactly half the volume of the mother. Each mRNA, immature GFP, and mature GFP in the mother cell is assigned to one of the daughters with equal probability. The simulation subsequently tracks only one of the daughters (which is randomly chosen).

4. During the growth of each cell, the following stochastic events are simulated using the Gillespie algorithm [65]:

- A new instance of DNA damage is created at rate *r*_*b*_.
- Each existing DNA damage instance is repaired at rate *r*_*r*_.
- If the promoter is not bound by LexA, LexA binds to the promoter at rate proportional to the current concentration of LexA. Note that, when LexA is at its equilibrium concentration in the absence of DNA damage, this rate is *L*_SS_.
- If the promoter is bound by LexA, LexA unbinds at a fixed rate *L*_off_, which is determined by the strength(s) of the LexA binding site(s) in the promoter.
- If the promoter not bound by LexA, an new mRNA molecule is created (i.e. transcribed) at rate *r*_*m*_*V*. Note that we assume that the transcription rate is proportional to cell volume *V* because plasmid copy number is approximately proportional to cell volume and we assume that the concentration of mRNA polymerase is approximately constant across the cell cycle. That is, we assign larger cells a larger transcription rate, but do not explicitly model the transcription dynamics at the individual plasmids.
- Each molecule of mRNA decays at a rate *δ*_*m*_.
- Each molecule of mRNA is translated into immature GFP at rate *r*_*p*_.
- Each immature GFP matures into fluorescing GFP at rate *r*_*g*_.
- Each mature GFP disappears to due bleaching (or decay) at a total rate *δ*_*g*_.

Note that, in the above simulation algorithm, growth and division of the cells are deterministic whereas all gene expression processes are stochastic.

The concentration dynamics of LexA follows one of the following two scenarios. In the equilibrium model, the concentration of free LexA in the cell is assumed to be constant and dependent on the amount of Ciprofloxacin (i.e. there is a fixed LexA concentration at each Ciprofloxacin level). In the dynamic model, the concentration of LexA in the absence of DNA damage is assumed constant and at a level corresponding to a binding rate *L*_SS_. Whenever DNA damage is present, the concentration is assumed to decay exponentially at a rate *α*. After the damage is repaired, we assume that the concentration of LexA recovers at a rate *β* until it saturates back to the steady-state rate *L*_SS_. We generally assume for simplicity that *α* and *β* are equal, i.e. that it takes roughly as much time for LexA concentration to decrease as it takes to recover after the DNA damage is repaired. Note that, while the introduction and repair of instances of DNA damage is stochastic, the LexA concentration in responses to these DNA damage instances is deterministic.

## 6 Theoretical analysis of the dynamic model

In this section we present a number of analytical derivations to characterize how peak durations, heights and frequencies are related to the parameters of the dynamic model (Table 1 in the main text). In particular, we will show that, in order to fit the experimental data, instances of DNA damage must be repaired relatively quickly, that in this regime peak heights are exponentially distributed, and that the ratio of the promoter’s transcription initiation rate *r*_*m*_ and the rate of DNA damage repair *r*_*r*_ sets the average peak height.

### 6.1 DNA damage events must be short-lived

The duration of the peaks is determined by different time scales, including the DNA damage repair rate *r*_*r*_, the decay rate *δ*_*m*_ of the mRNA, and the maturation rate *r*_*g*_ of the GFP. Here we first show that, because the GFP maturation time is known to be 6 − 8 minutes [53, 54] and the mRNA decay times are typically in the range of 2 − 10 minutes [61], even an instantaneous burst of mRNA production would lead to a peak in volumic production *q*(*t*) of a duration that is only a little bit less than the observed durations of the peaks (which have an approximately Gaussian distributed duration with mean around 8 minutes).

Let’s assume that at time *t* = 0 an instantaneous burst of *m*_0_ mRNA molecules is produced, which then decay at a rate *δ*_*m*_ and are translated into immature GFP proteins at a rate *r*_*p*_. In addition, immature GFP proteins mature into fluorescing GFP at a rate *r*_*g*_. The volumic production rate *q* is defined as the production of mature GFP per unit time and volume.

For this simple analytical argument, we will ignore stochastic fluctuations and treat mRNA number as a continuous quantity that simply decays exponentially with time

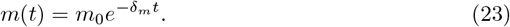

Similarly, for the total amount of immature protein *P* we write

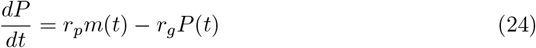

and the volumic production is related to *P* (*t*) through

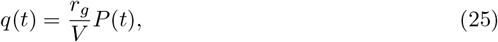

where *V* is the volume of the cell. Note that since the peak durations are short compared to the cell cycle, we will ignore the growth of the volume *V* during the peak. Solving for *q*(*t*) we find

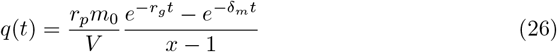

where 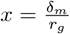 is the ratio of the mRNA decay to the protein maturation rate.

Let *t*^*^ denote the time at which the maximum of the peak in volumic GFP production *q*(*t*) occurs, *q*^*^ the height of this maximum, and *dt* the time for which the peak is above half of its maximum. First, the maximum production occurs at

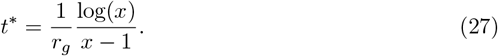

Using this, we find for the height of the peak

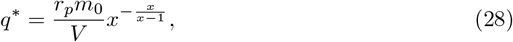

which is proportional to the mRNA burst size *m*_0_ and inversely proportional to the volume *V*. Note, however, that below we will see that this dependence on volume will cancel out since the mRNA burst sizes *m*_0_ are themselves proportional to volume *V*.

Finally, there is no simple analytical expression for the time *dt* that *q*(*t*) is larger than half its maximum *q*^*^*/*2. However, from equation (26) we numerically find that with a GFP maturation rate of *r*_*g*_ = 1*/*7 minute, *dt* runs from *dt* ≈ 11 minutes at a *δ*_*m*_ = 1*/*2 per minute mRNA decay rate to *dt* ≈ 18 minutes at *δ*_*m*_ = 1*/*6 per minute. The latter is already longer than the width of the peaks we observe in the data (full width half maximum: 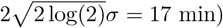). That, using the known GFP maturation time of about 7 minutes, we have to assume mRNA lifetime is less than 6 minutes in order for the peak duration from an instantaneous burst of mRNA production to remain below the experimentally observed peak durations. Consequently, the mRNA production bursts most be short, i.e. at most a few minutes long.

The parameter that controls the duration of transcriptional bursts is the DNA damage repair rate *r*_*r*_, i.e. the average duration of DNA damage events is 1*/r*_*r*_ minutes. For a relatively high repair rate of *r*_*r*_ = 0.8 per minute, only one in fifty DNA damage events take longer than 5 minutes to repair, and we used our simulation framework to compare the peak durations at this repair rate with those observed experimentally as we varied mRNA lifetime from 1.5 − 4.5 minutes and GFP maturation time ranging from − 8 minutes.

Supplementary Figure S11A shows that, with this repair rate, the mean duration of the peaks matches the experimentally observed mean duration for maturation times of 7 − 8 minutes when mRNA lifetime is 3.5 − 4.5 minutes. Moreover, at an mRNA decay time of 4 minutes and a maturation time of 7 minutes, the distribution of durations of the simulated peaks exactly matches those observed in the experiments when the repair rate is *r*_*r*_ = 0.8 per minute (Suppl. Fig. S11B).

### 6.2 When damage events are short-lived, peak heights are exponentially distributed

As we have seen in the last section, the short durations of the observed peaks in volumic production require that instances of DNA damage are repaired relatively quickly. We also saw that the observed distribution of peak durations can be well fit by the model when the repair rate is *r*_*r*_ = 0.8 per minute. We first used our simulation framework to investigate whether the distribution of peak heights also matches the exponential distribution that is observed experimentally. We find that, for a relatively narrow range of repair rates from *r*_*r*_ = 0.5 to *r*_*r*_ = 1.0 per minute, the distribution of peak heights in the simulations indeed matches the experimental distribution observed experimentally (Suppl. Fig. S12). However, from repair rates of *r*_*r*_ = 0.4 per minute or below, and from *r*_*r*_ = 1.5 per minute or above, the distributions of peak heights already start deviating from what is observed experimentally (Suppl. Fig. S12).

**Fig S11.**
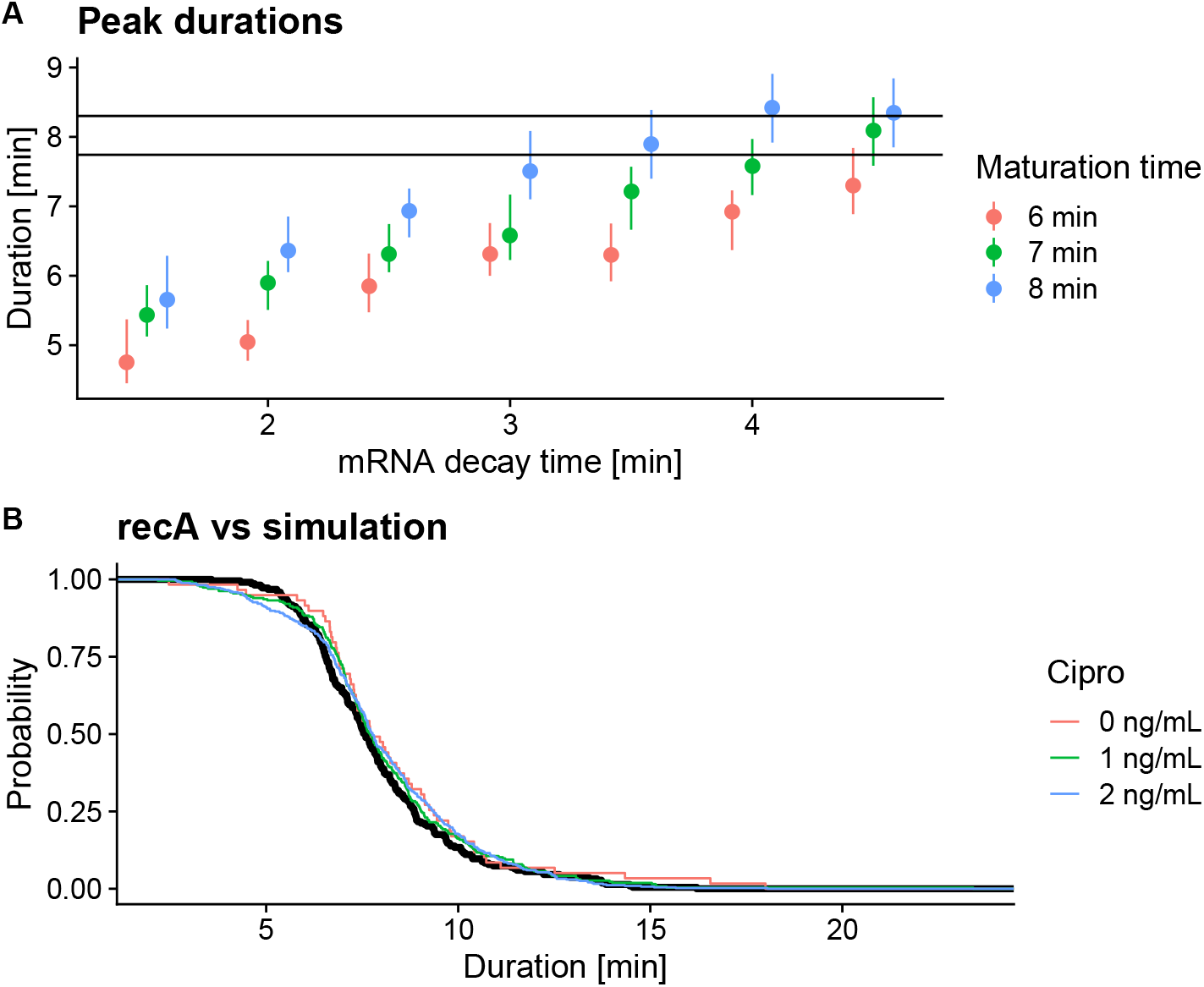
Duration of peaks in volumic GFP production from bursts of mRNA production. **A**: Median peak durations (mean ± standard-error of the standard-deviations of the Gaussians fitted to peaks in volumic production) in simulations of the dynamic model using different values for the mRNA decay rate (horizontal axis) and the GFP maturation time (symbol colors, see legend) and a DNA damage repair rate of *r*_*r*_ = 0.8min^−1^. The black horizontal line corresponds to the average duration of the peaks in the experimental data for recA. **B**: Reverse cumulative distributions of peak durations for the simulations of the dynamic model (black) and for the experimental data from the recA promoter in different conditions (colored lines). For this simulation mRNA decay time was 4 minutes, GFP maturation time was 7 minutes and repair rate again *r*_*r*_ = 0.8min^−1^. Note that not only the median but also the variation in peak durations matches between experiments and simulations.

**Fig S12.**
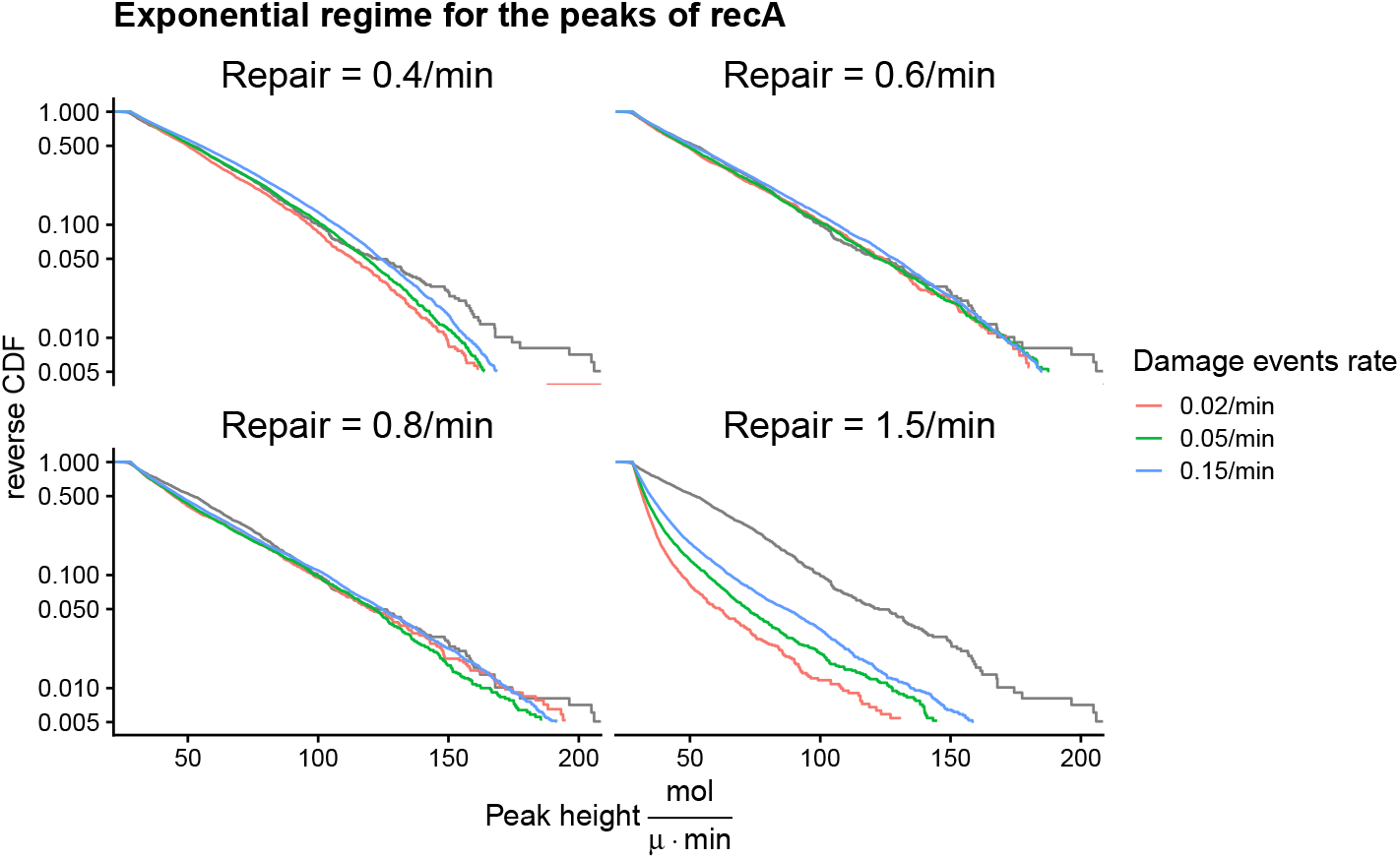
Reverse cumulative distributions of peak heights from simulations of the dynamic model. Each panel shows the reverse-cumulative distributions of peak heights obtained using simulations of the dynamic model with the recA promoter for a given repair rate (shown at the top of the panel) and for different frequencies of DNA damage events (different colored lines, see legend). The transcription rate *r*_*m*_ has been set so as to match the average of the exponential distributions observed experimentally (black lines). Vertical axes are shown on a logarithmic scale. Note that at the lowest and highest of these repair rates the distributions of peak heights starts deviating from a pure exponential.

Here we provide a simple analytical argument to explain why, when DNA damage events are short-lived, the distribution of peak heights has an exponential tail.

Equation (28) shows that, when mRNAs are produced in a short burst, the height of the resulting peak in volumic production is proportional to the number of mRNAs *m* produced in the burst, and inversely proportional to the volume *V* at which the burst occurred. In order for a transcriptional burst to occur, two conditions need to be satisfied. First, LexA needs to unbind from the promoter and second, the LexA concentration needs to be low enough such that the transcription rate *r*_*m*_*V* is at least on the order of the LexA rebinding rate, which only occurs during DNA damage events. Note that the longer the DNA damage lasts, the lower the LexA concentration and the lower the rate of LexA rebinding. Thus, the larger bursts of mRNA production correspond to DNA damage events that last longer. The exponential tail of the distribution of peak heights thus corresponds to relatively long DNA damage events where the LexA concentration has become low.

For these longer DNA damage events, we will approximate the LexA concentration dynamics by ignoring the transient phase during which the concentration drops from *L*_ss_ to a new low steady-state at *L*_low_, as well as the transient recovery phase during which it grows from *L*_low_ back to *L*_ss_. That is, we model such DNA damage events as if the LexA concentration is *L*_low_ during the entire DNA damage event.

During such DNA damage events the following events can occur stochastically: LexA-bound promoters switch to unbound at rate *L*_off_, unbound promoters switch to LexA-bound at rate *L*_low_, new transcripts are initiated from unbound promoters at rate *r*_*m*_*V*, and the damage event ends at a rate *r*_*r*_. Because these rates are constant, it is straight-forward to calculate the probability *P*_*m*_ that *m* mRNAs have been created at the time that the damage event ends.

At any time, the promoter can be either in the LexA-bound state *b* or unbound state *u*. If the promoter is in state *b*, only two events can ocur next. Either the promoter switches to unbound, or the DNA damage event ends. The probability *P* (*u*| *b*) that the next event is a switch to the unbound state is

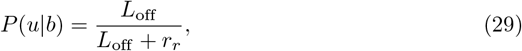

and the probability *P* (*e*|*b*) that the next event is the end of the damage event is

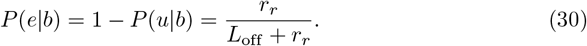

If the promoter is in the unbound state *u* the possible next events are a switch to the bound state *b*, the end of the damage event, or the generation of a new transcript. The probability that the next event is a switch to the bound state is

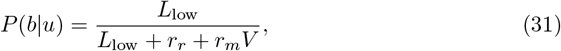

the probability that the next event is the end of the damage event is

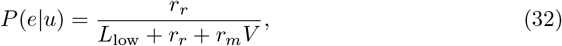

and the probability *P* (*t*|*u*) that the next event is the initiation of a transcript is

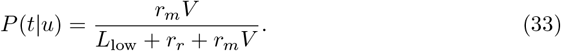

Imagine that the promoter is in state *u* and that *m* mRNAs have already been generated. We now want to calculate the probability *ρ* that, before the damage is repaired, another transcript is generated. This can happen either by directly generating a transcript with probability *P* (*t* |*u*), or switching back and forth from bound to unbound and back *k* times with probability [*P* (*b*|*u*)*P* (*u*|*b*)]^*k*^ before creating the transcript. We thus have

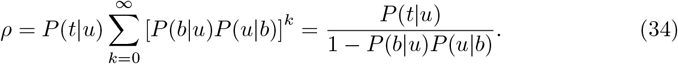

Thus, given that the promoter starts out in the unbound state, with probability *ρ* another transcript is created, and with probability 1 − *ρ* the damage event ends.

Thus, assuming that at the start of the damage event the promoter is bound, the probability that 0 transcripts are generated at the time the damage is repaired is

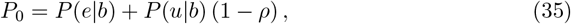

and the probability that *m* ≥ 1 transcripts are generated is

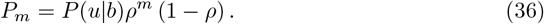

Note that this is a geometric distribution which is the discrete equivalent of the exponential distribution. The average number of mRNAs produced in the burst is given by

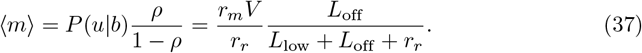

Note that equation has an intuitive interpretation. The first factor *r*_*m*_*V/r*_*r*_ is the expected number of transcripts that would be produced if the promoter were unbound during the entire event. The second factor is the expected fraction of the time that the promoter is unbound during the event, which depends on the strength of the binding site.

Therefore, using equation (28), the distribution of peak heights *q*^*^ resulting from bursts that occurred at volume *V* is approximately exponential with mean

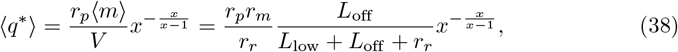

which, notably, is independent of *V*, so that one exponential describes peaks resulting from bursts occurring at different cell volumes *V*.

In summary, the tail of the distribution of peak heights is approximately exponentially distributed with a mean that is proportional to transcription rate *r*_*m*_ of the promoter. Thus, the our dynamical model can be made to fit the exponential height distributions of each promoter by simply setting the transcription rate *r*_*m*_ so as to fit the average peak heights.

However, it should be noted that this derivation relied on the assumptions that damage events are short-lived. As Suppl. Fig. S12 shows, at a repair rate of 0.4/min the DNA damage events start to be sufficiently long that the assumption that transcriptional bursts are short starts breaking down, and we notice that the height distribution starts to bend away from a pure exponential tail. We also see that at very high repair rates (i.e. *r*_*r*_ = 1.5 per minute), longer damage events become so rare that the exponential tail no longer holds over the range of peak heights that we observed in experiments. That is, apart from the burst durations, also the fact that the observed distribution of peak heights is exponential restricts the repair rate to lie in a relatively narrow range.

In Suppl. Fig. S13 we show that for repair rates in this range, we can also recapitulate the observed absolute frequencies of peaks as a function of cut-off *q*_*c*_ for recA, provided that the repair rate lies within the relatively narrow range of 0.5 − 1.0 per minute.

**Fig S13.**
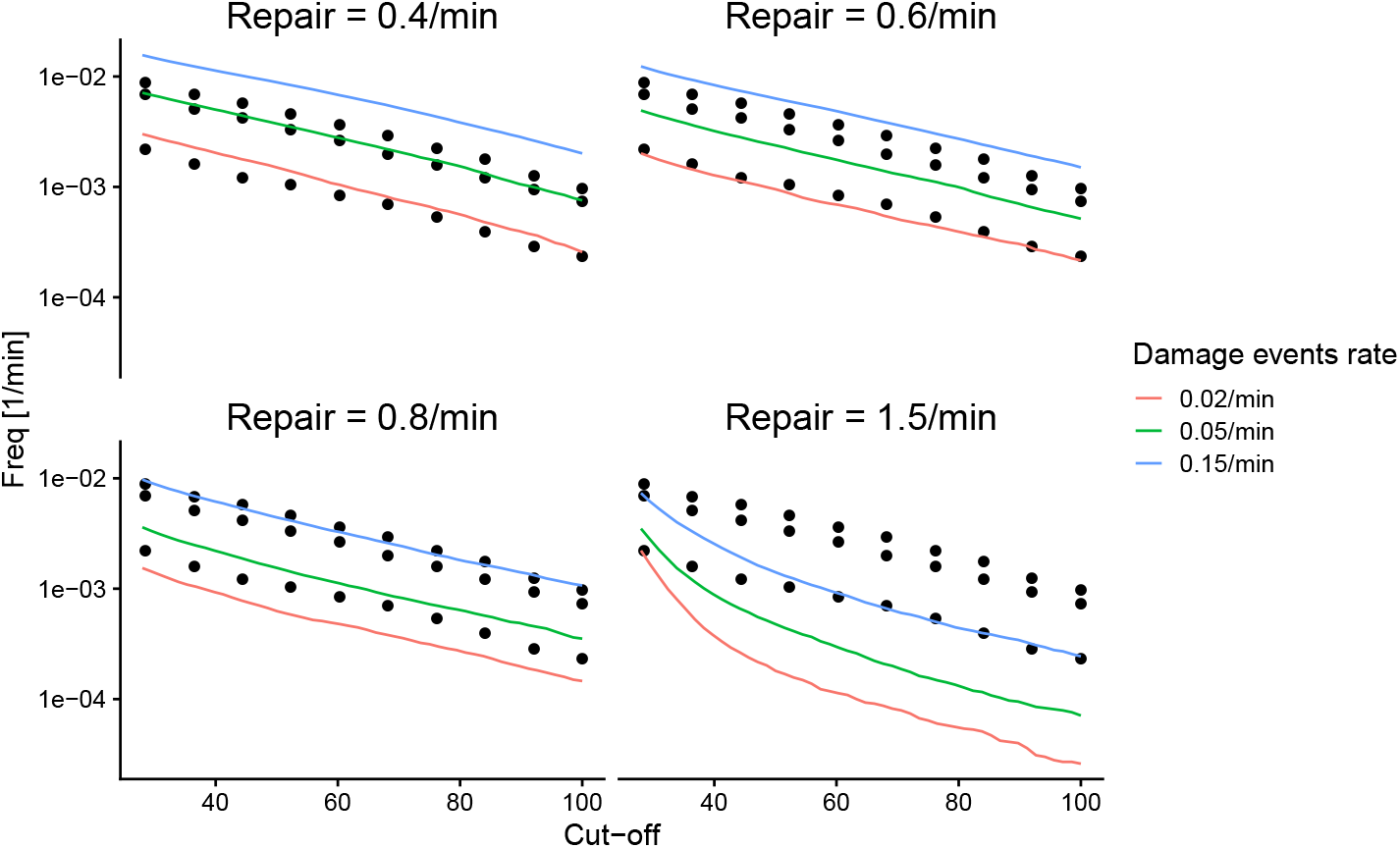
Peak frequencies in simulations of the dynamical model with the recA promoter. Each panel shows the observed peak frequenciwa as a function of cut-off in peak height (horizontal axis) from simulations of the dynamical model with the recA promoter for different frequencies of DNA damage events (colored lines, see legend) and for a given rate of damage repair (indicated at the top). For reference, the black dots show the frequencies observed experimentally at different Cipro concentrations (i.e. the same data as in Fig. S8). The vertical axes are shown on a logarithmic scale. The results show that, by appropriately setting the frequency of DNA damage events in the model, the peak frequencies as a function of cut-off can be matched between simulations and experiments, provided that the repair rate lies in a relatively narrow range of 0.5 − 1.0 per minute.

Finally, although our simple model ignores relevant fluctuations such as fluctuations in growth rate and the unequal sizes of daughters at division, Fig. S14 shows that it is able to capture the distributions of GFP concentrations across cells and Cipro concentrations for the recA promoter with reasonable accuracy.

**Fig S14.**
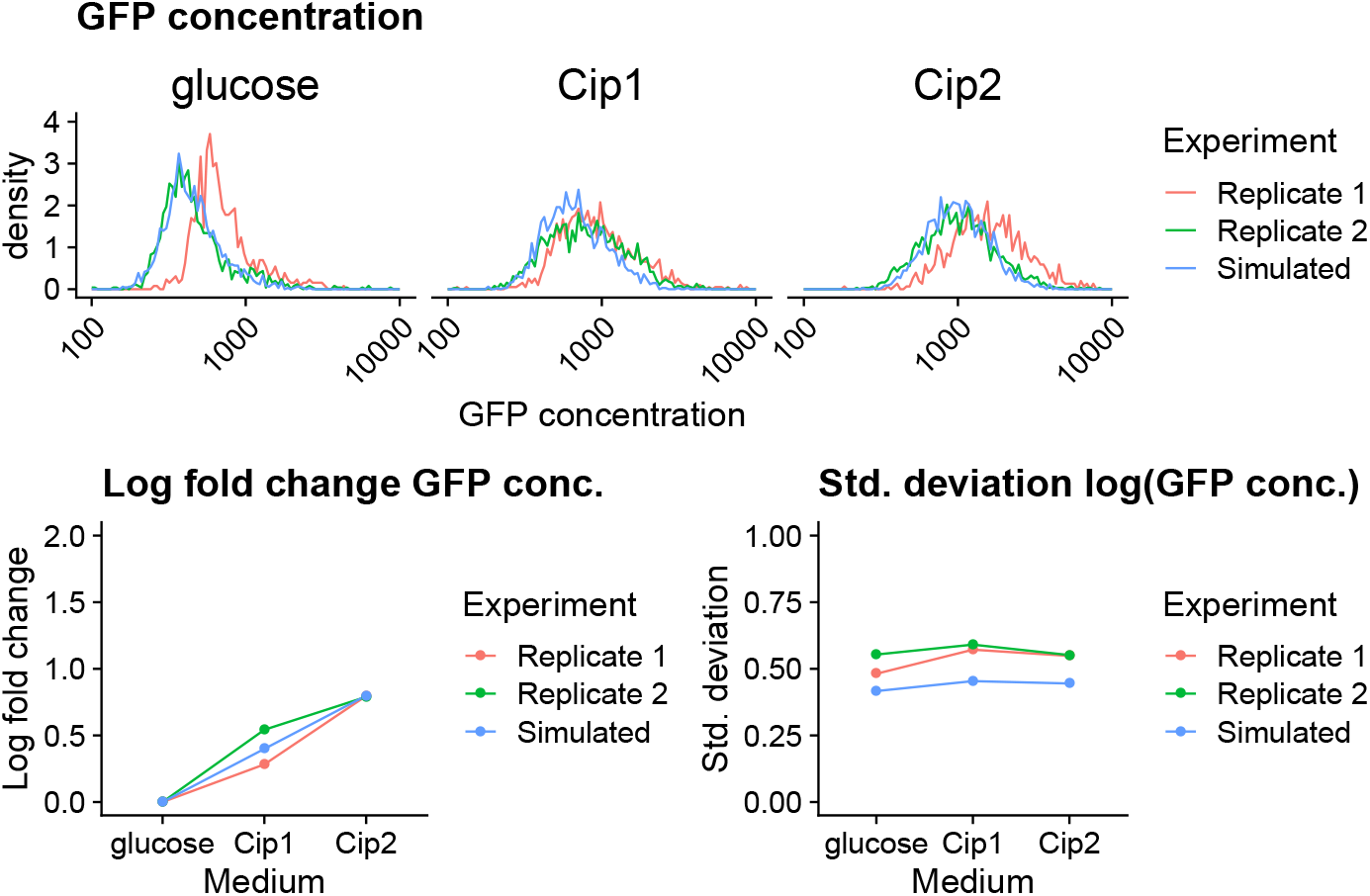
Distribution of simulated vs observed GFP concentrations. **Top**: Comparison between the GFP concentration distributions for the recA promoter as observed in the two replicates and in the simulations of our simple dynamical model (colors) for each of the Cipro concentrations (panels). **Bottom left**: Log-fold changes in the average expression of the recA promoter as a function of Cipro concentration for the two replicates and the simulations. **Bottom right**: Standard-deviation in expression of the recA promters as a function of Cipro concentration for the two replicates and the simulations. Note that the model fairly accurate captures the GFP concentration distributions and fold-changes observed in the experiments although it somewhat underestimates the standard-deviation across cells.

